# ErbB2/HER2 governs CDK4 inhibitor sensitivity and timing and irreversibility of G1/S transition by altering c-Myc and cyclin D function

**DOI:** 10.1101/2024.05.09.593450

**Authors:** Ayaka Nagasato-Ichikawa, Ken Murakami, Kazuhiro Aoki, Mariko Okada

## Abstract

Cell cycle entry and the irreversible transition from the G1 to S phase are crucial for mammalian cell proliferation. Among the ErbB family, the ErbB2/HER2 receptor is a key driver of cancer growth. However, the quantitative mechanisms underlying the ErbB2-mediated G1/S transition remain unclarified. Here, we performed an extensive time-course analysis of high and low ErbB2-expressing breast cancer cells to describe the regulatory mechanisms of the ErbB2-mediated G1/S transition. Live-cell imaging using cell cycle reporters revealed that the G1/S transition occurs 20 h after ErbB2 activation, driven primarily by the cyclin D1/CDK4–RB axis. Hsp90 is regulated by CDK4 activity and controls the stability of ErbB2 protein in a time-dependent manner. CDK4 inhibitor treatment arrested the cell cycle in most cells; a subpopulation showed a 25-h delay in G1/S entry associated with enhanced c-Myc activation. In high ErbB2-expressing cells, CDK4 inhibition led to c-Myc overactivation, a rapid decrease in cyclin D1 expression, and cell cycle arrest. Overall, we demonstrate how ErbB2 receptor levels modulate the roles of cyclin D1 and c-Myc in the G1/S transition and suggest that variations in ErbB2 levels within breast cancer tissues confer heterogeneous sensitivity to CDK4 inhibitors, potentially complicating treatment.

## Introduction

The initiation of the cell cycle and subsequent transition from the G1 to S phase are regulated by a complex molecular network involving the formation of a cyclin D1/CDK4 protein complex (*1–3*). Understanding the regulation of the G1/S transition by mitogenic signals is fundamental for comprehending cell biology and developing cancer treatments that overcome drug resistance. For example, based on the important role of cyclin D1 overexpression for enhancing breast cancer cell proliferation (*4*), various cell cycle inhibitors targeting CDK4 have been developed to treat breast cancer (*5*, *6*).

Breast cancer can be classified into four subtypes based on the expression of estrogen receptors (ERs), progesterone receptors, and ErbB2 (HER2) receptors: luminal A (ER^+^ and HER2^-^), luminal B (ER^+^ and HER2^+^), ErbB2/HER2-positive (ER^-^ and HER2^+^), and triple-negative (ER^-^ and HER2^-^) (*7–9*). CDK4 inhibitors (CDK4is) have been used to treat the luminal A subtype (*5*, *10*), but long-term administration has led to drug resistance (*11–17*). ErbB2 tyrosine kinase activity, commonly elevated in malignant breast cancer, increases cyclin D1 levels (*18*, *19*). Stimulation of ErbB2 receptors activates the MAPK/ERK and PI3K/AKT signaling pathways, leading to the rapid induction of early-response genes such as *c-fos* and *c-myc* (*20–22*)—key regulators of cyclin D1 and CDK4 expression (*22–24*). Both c-Myc and cyclin D1 are involved in the G1/S transition, though they appear to have distinct roles (*25*). Therefore, we speculated that the small population of high ErbB2-expressing cells in the luminal A subtype may contribute to CDK4i resistance by rewiring the G1/S network during the early stages of drug treatment. However, the molecular mechanisms by which ErbB2 affects the G1/S transition following CDK4i treatment remain unclear.

The G1/S transition typically involves signal-dependent and -independent processes, which are distinguished by the restriction (R) point (*2*, *26*). The mitogenic signal primarily regulates the expression and activation of cyclin D1/CDK4, which phosphorylates retinoblastoma protein (RB) prior to the R point. Thus, this signal-dependent process is primarily driven by the cyclin D1/CDK4-RB axis (*2*). The phosphorylation of RB (p-RB) activates E2F1, promoting the expression of its target genes, including cyclin E. Cyclin E then forms a complex with CDK2, which further phosphorylates p-RB, resulting in a fully phosphorylated state (pp-RB). Fully activated pp-RB enhances E2F1 activity, creating a mitogen-independent cyclin E/CDK2–RB–E2F1 positive feedback loop (*27*, *28*). However, the expression and stability of c-Myc are modulated by external signals (*29*), and c-Myc directly influences E2F1 promoter activity (*30*). Consequently, the core machinery of cyclin E/CDK2–RB–E2F1 is considered to be indirectly regulated by mitogenic signals.

ERK controls the expression of c-Myc and cyclin D1 (*31*, *32*), and AKT regulates their protein stability (*31*, *33*). However, p27 expression, which plays a crucial role in the accumulation of cyclin D1 and CDK4 during the G1 phase (*34*), is repressed by c-Myc (*35*). Since the ErbB2 receptor activates the ERK and AKT pathways at different magnitudes (*36*), different levels of ErbB2 may modulate the core machinery of the G1/S network and, consequently, the timing of cell cycle entry through modulation of cyclin D1 and c-Myc.

Therefore, investigating the ratio of c-Myc to cyclin D1 at the single-cell level may elucidate their specific functions in the G1/S transition and the regulatory role of the ErbB2 receptor. In the present study, we used representative luminal A subtype cell lines (MCF-7 and T47D), which express low levels of the ErbB1 receptor (EGFR) and high levels of the ErbB3 receptor (*22*, *36*, *37*). ErbB2 is an orphan receptor that is activated by heterodimerization with ErbB3, ErbB4, or EGFR (*21*). Stimulation with the ErbB3 ligand, heregulin (HRG), triggers the heterodimerization of ErbB2 and ErbB3, thereby activating the ERK and AKT pathways (*36*). We conducted time-course protein phosphorylation profiling after HRG stimulation, fluorescent ubiquitination-based cell cycle indicator (FUCCI) live-cell imaging for cell cycle dynamics, and transcriptomic/epigenetic studies to identify key transcription factors during CDK4 inhibition. We evaluated the role of ErbB2 in regulating the timing and irreversibility of the G1/S transition, revealing the molecular mechanism underlying heterogeneous CDK4 inhibitor sensitivity.

## Results

### G1/S transition occurs approximately 20 h after HRG stimulation

First, we used FUCCI (*38*) live-cell imaging to analyze the cell cycle dynamics in MCF-7 or T47D cells after HRG stimulation (Fig. 1A) and analyzed the distribution of each cell cycle phase in the presence or absence of HRG for up to 72 h in MCF-7 cells (Fig. 1B–D). HRG stimulation began to significantly increase the proportion of S/G2/M-phase cells after 24 h compared to the control. The number of nuclei, serving as a proxy of cell number, increased 1.4-fold after 48 h, suggesting that HRG stimulation enhances cell proliferation. Conversely, control cells showed a gradual increase in the proportion of S/G2/M cells, but there was no increase in the number of nuclei, suggesting cell cycle arrest during the S/G2/M phase. To further examine the times of transition for each cell cycle phase, we performed single-cell tracking and quantified the distribution of the cell cycle phases (Fig. 1E; Fig. S1A, B). The G1/S transition occurred at 19.0 ± 7.2 h after HRG stimulation. In comparison, the S/G2 and M/G1 transitions occurred at 32.5 ± 9.3 h and 39.4 ± 8.7 h, respectively (Fig. 1F). These findings suggested that most cells entered the S phase approximately 20 h after HRG stimulation, with some intercellular variability.

**Fig. 1.**
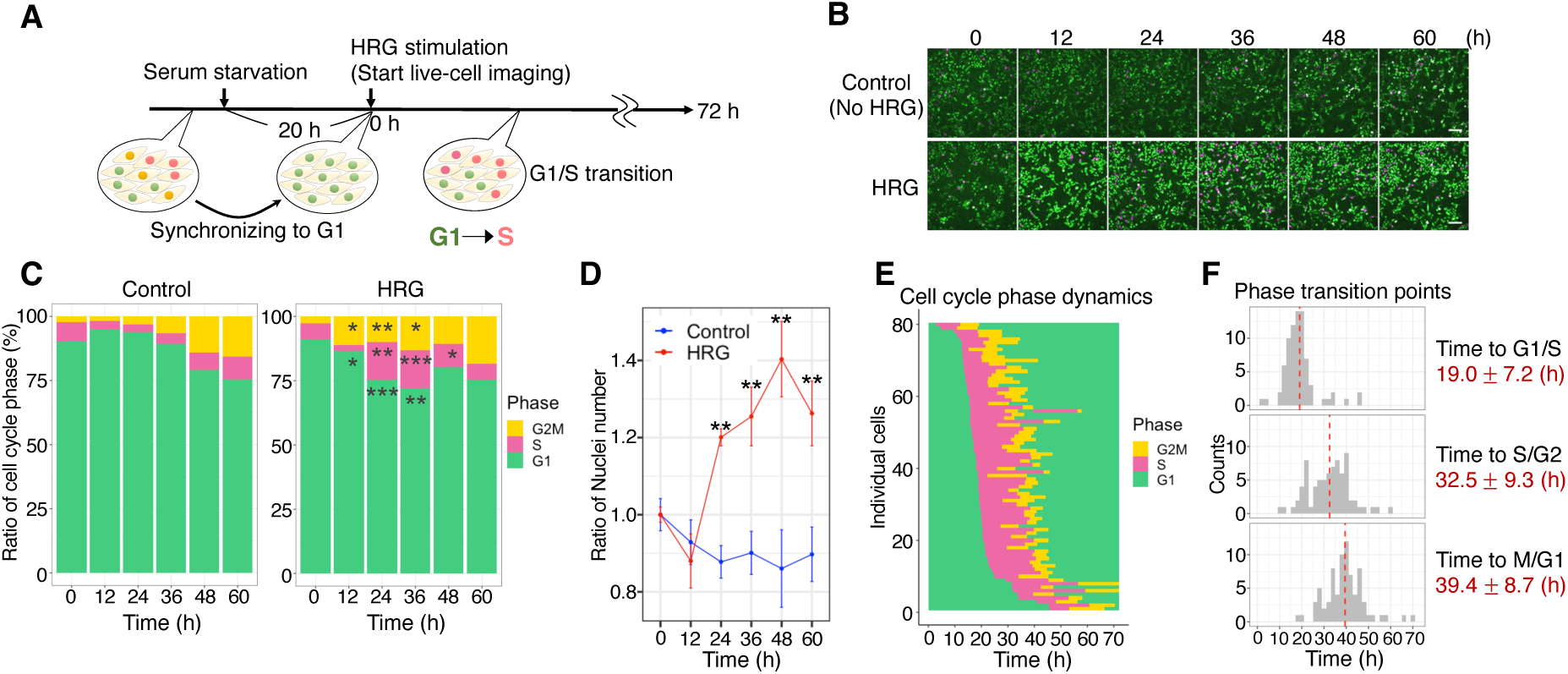
G1/S transition occurs approximately 20 h after HRG stimulation. (A–F) MCF-7 cells stably expressing FUCCI. (**A**) Experimental scheme for live-cell imaging using FUCCI biosensor. (**B**) Images were captured every 20 min up to 72 h in conditions with or without 10 nM HRG. Scale bar: 100 µm. (**C**) Percentage distribution of each cell cycle phase (%) in each image; *n* = 3 fields per image. (**D**) The number of nuclei increased with each passing hour relative to the cell number at 0 h. (**E**) Cell cycle phase dynamics estimated based on Geminin and PIP fluorescence for individual cells followed up to 72 h; *n* = 79. (**F**) Histogram of cell cycle transition times for cells that completed the cell cycle within 72 h (*n* = 77), based on the results in (E). Red dotted lines indicate the mean ± SE of each transition time. In (C) and (D), \**P* < 0.05, \*\**P* < 0.01, \*\*\**P* < 0.001 (with respect to control and HRG; *t*-test).

### CDK4 and cyclin D1 proteins are expressed in a time-dependent manner at the G1/S transition

We further examined the signaling dynamics of the ErbB2, AKT, and ERK pathways (*21*, *39*) following HRG stimulation (Fig. 2A, B). p-ErbB2, p-AKT, and p-ERK levels decreased after peaking within 30 min, and remained present until the S/G2 transition point (32.5 ± 9.3 h). Although c-Fos protein levels peaked within 2 h and quickly reduced to basal levels, p-c-Fos showed similar dynamics to those of p-ERK, a critical regulator of p-c-Fos (*22*). Following ErbB2 activation, p-AKT, p-ERK, and p-c-Fos levels remained consistent whereas c-Myc and p-c-Myc (Ser^62^) levels decreased around the time of the G1/S transition (19.0 ± 7.2 h). We estimated the timing of the R point based on the dynamics of cell cycle regulatory proteins (Fig. 2C, D; Fig. S2A, B). The protein levels of cyclin D1 and CDK4 started to increase shortly after HRG stimulation, peaked at approximately 20 h (G1/S transition point), and then remained consistent until the S/G2 transition (Fig. 2D), consistent with the trend in p-RB (Thr^373^) expression (*40*). p-RB (Ser^807/811^) showed a monotonic increase similar to cyclin E (Fig. S2A, B), suggesting the involvement of CDK2-mediated S phase progression (Fig. S2A, B). Since the R point is found just before the G1/S transition, we estimated that it occurs at 12–16 h, when the E2F1 protein level is highly elevated. Thus, our results substantiate that cyclin D1/CDK4 maintains RB phosphorylation near the R point, which is a critical initial step toward the G1/S transition associated with HRG-induced ErbB2 activation.

**Fig. 2.**
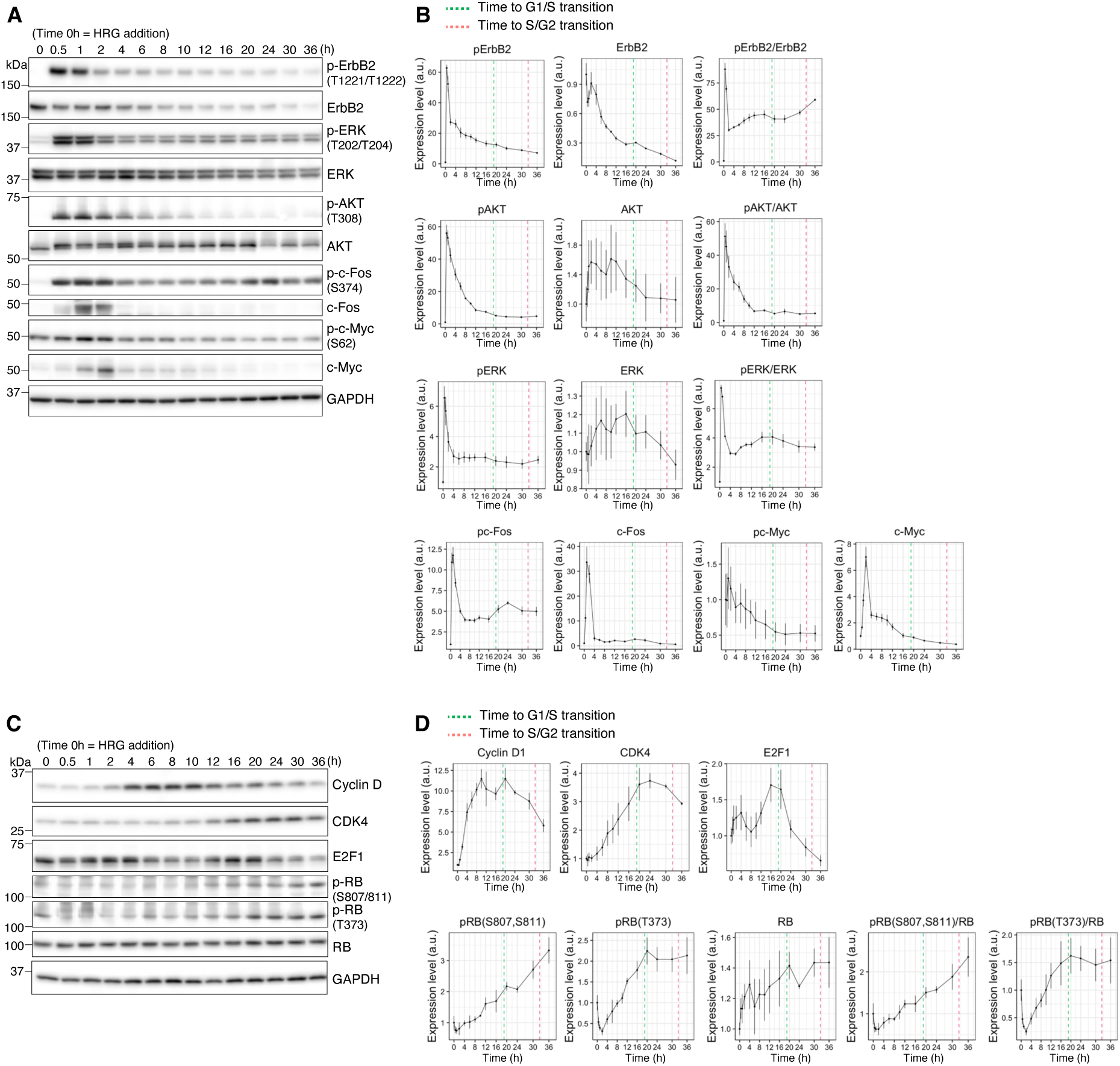
Time-dependent expression of CDK4 and cyclin D1 in G1/S transition. (**A–D**) Dynamic changes in the expression or phosphorylation of individual proteins in MCF-7 cells stably expressing FUCCI, detected by western blotting up to 36 h after treatment with 10 nM HRG. (**A, B**) ErbB2 pathway proteins. (**C, D**) Proteins regulating G1/S transition. Protein and phosphorylation levels were normalized relative to those of GAPDH; *n* = 3.

### A population subset enters the cell cycle even after CDK4 inhibition

We confirmed the involvement of cyclin D1/CDK4 in the HRG-stimulated G1/S transition. Cells were treated with CDK4i at 12 h after HRG stimulation (before the R point; Fig. S3A, B). As ERK activates cyclin D1 expression and AKT inhibits its degradation (*41*), we used AKT inhibitor (AKTi) and MEK inhibitor (MEKi), which have comparable inhibitory effects on cyclin D1 expression (Fig. S3A, B), as CDK4is. We examined the proportions of cells in each cell cycle phase at 32 h and 72 h after HRG stimulation (Fig. 3A, B). At 32 h, fewer cells treated with CDK4i (13.8%), AKTi (7.05%), or MEKi (13.8%) were in the S/G2/M phase compared to non-treated cells (29.2%). At 72 h, AKTi (9.85%) and MEKi (5.50%) reduced the proportion of cells in the S/G2/M phase, but CDK4i (17.7%) did not. At this time point, the number of nuclei was significantly reduced by AKTi and MEKi but not CDK4i (Fig. 3C). This finding indicated that AKTi and MEKi induced the growth of most cells, whereas a significant subpopulation of cells were not affected by CDK4i, suggesting that additional mechanisms maintain cell cycle functionality in the absence of CDK4.

**Fig. 3.**
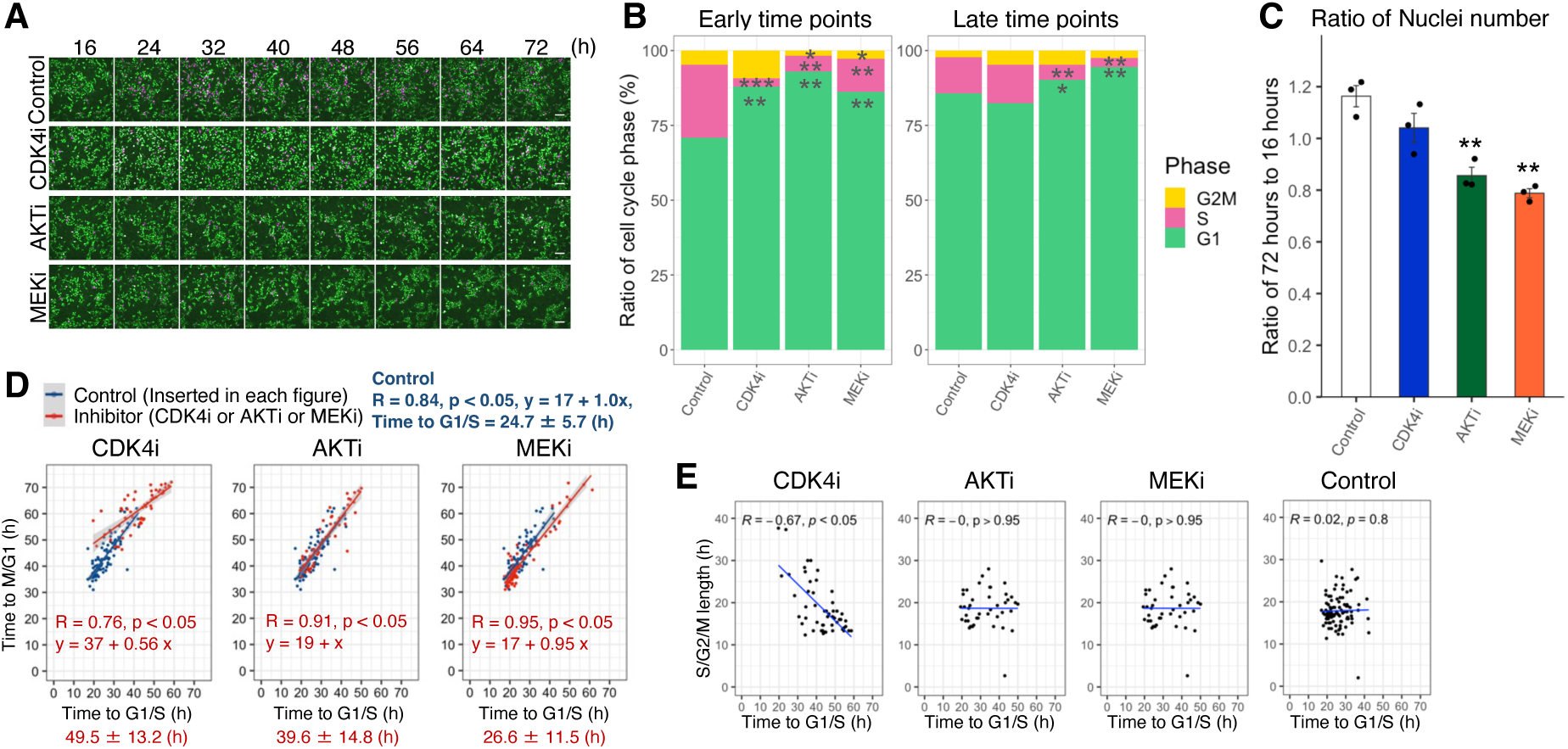
A population subset enters the cell cycle even after CDK4 inhibition. (**A**) Each inhibitor (250 nM CDK4i; 600 nM AKTi; 500 nM MEKi) was used to treat the FUCCI stably expressing MCF-7 cells 12 h after HRG stimulation. Images were acquired every 20 min from 16 to 72 h after HRG stimulation. Scale bar: 100 µm. (**B**) Percentage distribution of each cell cycle phase (%) in the overall image at 32 h and 72 h after HRG stimulation; *n* = 3. (**C**) Increasing number of nuclei from 16 h to 72 h after HRG stimulation; *n* = 3. (**D**) Data from cells that completed the M/G1 transition within 72 h. Control, *n* = 96; CDK4i, *n* = 52; AKTi, *n* = 42; MEKi, *n* = 75. Times to G1/S (x-axis) and M/G1 (y-axis) transitions, with slope (*s*) and intercept (*i*) obtained from the regression equation. (**E**) Comparison of Pearson’s coefficients using data from (D). In (B) to (C), \**P* < 0.05, \*\**P* < 0.01, \*\*\**P* < 0.001 (with respect to control and each inhibitor; *t*-test).

To investigate the distinct mechanisms associated with CDK4i, we analyzed the correlations between the times to the G1/S and M/G1 transitions up to 72 h for the different treatments (Fig. 3D). Cells non-responsive to CDK4i showed significantly delayed G1/S transition times (49.5 ± 13.2) of ∼25 h compared to the control (24.7 ± 5.7), which tended to be longer than under AKTi (39.6 ± 14.8) and MEKi (26.6 ± 11.5; Fig. 3D). We analyzed the slope (*s*), representing the effect on the subsequent cell cycle duration after G1/S transition, and intercept (*i*), representing the time to the M/G1 transition, and observed a lower slope (*s* = 0.56) and higher intercept (*i* = 37) under CDK4i than under AKTi (*s* = 1.0, *i* = 19), MEKi (*s* = 0.95, *i* = 17), and control conditions (*s* = 1.0, *i* = 17). Thus, Pearson’s correlation coefficients (*R*) were significantly lower for CDK4i (0.76) than for AKTi (0.91), MEKi (0.95), and control conditions (0.84). This observation implied that the duration of the G1/S to G2/M phase might not be uniformly controlled only by CDK4i and phase-specific regulation might occur.

The CDK4i condition showed a clear negative correlation between the time to G1/S transition and duration of the S/G2/M phase (Fig. 3E). We divided the cell population into the early and late groups based on the median time of each G1/S transition (CDK4i, 49.5 ± 13.2; AKTi, 39.6 ± 14.8; MEKi, 26.6 ± 11.5; Control, 24.7 ± 5.7) and compared the duration of each S, G2, and M phase (Fig. S3C). The mean duration of S/G2/M differed significantly between the early (22.3 ± 7.2) and late (15.8 ± 2.4) groups, with a 6-h difference observed in the CDK4i groups. Thus, in this CDK4i-specific response, the majority of CDK4i non-responsive cells showed a long G1/S and short S/G2/M phase (Fig. 4A). We observed that CDK4i non-responsive T47D cells (another luminal A cell line), which normally express ErbB2 at a level three times higher than MCF-7 cells (Fig. S3D, E), also required a long time (32.6 ± 12.2 h) to reach the G1/S transition compared to the control (21.62 ± 6.2 h; Fig. S3F).

**Fig. 4.**
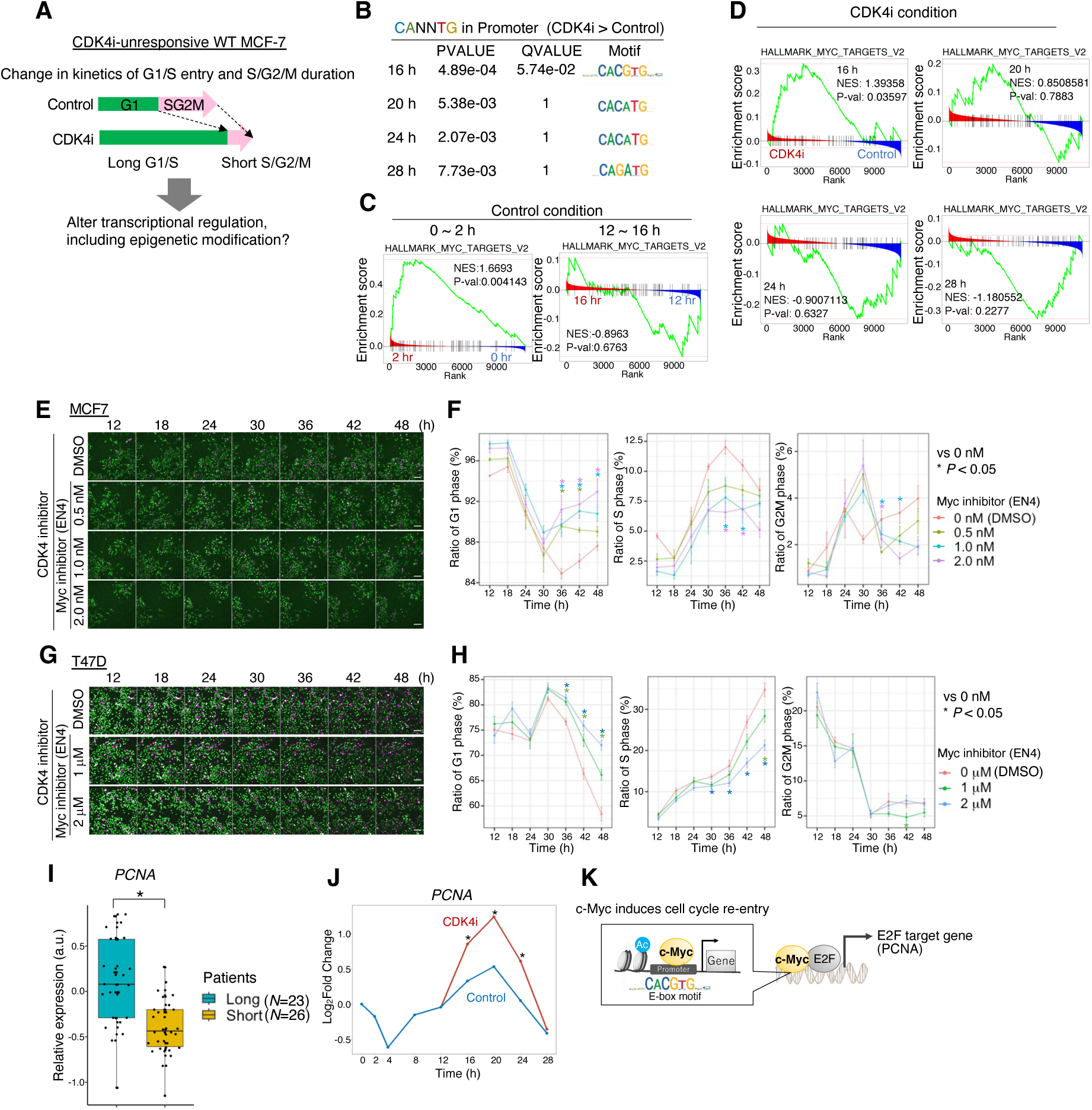
CDK4i enhances c-Myc transcriptional activity via H3K27Ac modification. (**A**) Changes in cell-cycle distribution following CDK4 inhibition. (B–H, J) MCF-7 cells stably expressing FUCCI. (**B**) Motif-enrichment analysis of H3K27Ac modifications in the promoter region. E-box sequences (CANNTG) were enriched in the promoter region at all time points in the CDK4i group compared to the control group. (**C**) GSEA was performed using the Myc-target gene set (*n* = 58) obtained from the Human MSigDB Collections. Comparison between 0–2 h after HRG stimulation (NES = 1.6693, *P* = 0.004143) and near the R point (12–16 h; NES = –0.8963, *P* = 0.6763). (**D**) GSEA was performed using the Myc-target gene set (C). Comparison between CDK4i and control conditions every 4 h (16 h: NES = 1.39358, *P* = 0.03597; 20 h: NES = 0.8508581, *P* = 0.7883; 24 h: NES = –0.9007113, *P* = 0.6327; 28 h: NES = –1.180552, *P* = 0.2277). (**E–H**) After stimulation with 10 nM HRG for 11.5 h, cells were treated with 0 (DMSO)/0.5/1.0/2.0 nM EN4 (E, F) or 0 (DMSO)/1/2 mM EN4 (G, H) for 30 min. Subsequently, MCF-7 (**E, F**) and T47D cells (**G, H**) were treated with 250 nM CDK4i. Scale bar: 100 µm. The percentage distribution of each cell cycle phase over time was quantified (**F, H**). (**I**) PCNA expression in microarray data from patients with luminal A subtype breast cancer undergoing long-term (16 weeks, *n* = 23) or short-term (4 weeks, *n* = 26) treatment with CDK4/6i (adj. *P* = 0.000132). (**J**) *PCNA* mRNA expression in RNA-seq data. (**K**) Increased c-Myc activity induces cell cycle re-entry to promote E2F target gene expression. In (F) and (H), \**P* < 0.05 (with respect to control and each inhibitor; *t*-test). In (I), \**P* < 0.05 (calculated using the Benjamini–Hochberg method). In (J), \**P* < 0.05 (calculated using EdgeR).

### CDK4i enhances c-Myc transcriptional activity via H3K27Ac modification

Given the distinct cell cycle kinetics observed under CDK4 inhibition (Fig. 4A), we performed transcriptome and epigenetic analyses to investigate the molecular mechanisms responsible for cell cycle regulation post CDK4 inhibition. To elucidate the changes in gene expression patterns, we added CDK4i 12 h after HRG stimulation and performed a time-course analysis of mRNA expression [0 (control), 2, 4, 8, 12, 16, 20, 24, and 28 h after HRG] as well as H3K27Ac and H3K4me1 modification [12 (control), 16, 20, 24, and 28 h after HRG]. First, active transcription factors were predicted at each time point with DoRothEA using RNA-seq data (*42*) (Fig. S4A). mRNAs corresponding to key transcription factors, such as *Myc*, AP-1 family (*Jun* and *Fos*), *EGR1*, *NFKB1*, and *STAT3*, which are known to promote cell cycle progression (*21*, *23*, *25*, *43*, *44*), were highly upregulated 16 h after HRG stimulation.

Motif-enrichment analysis was performed to confirm the binding of these transcription factors to the promoter regions. Here, we defined the regions without overlap between the H3K27Ac and H3K4me1 peak signals as the active promoter regions (*45*, *46*). E-box sequences (CANNTG) (*47*) were enriched in the promoter regions at all time points following CDK4i treatment (Fig. 4B; Fig. S4B, C). Given that c-Myc binds with E-box sequences (*48*), we focused on its role in G1/S transition. Control cells showed lower c-Myc protein and p-c-Myc (Ser^62^) levels near the R point (12–16 h after HRG) than during the early time point (0–2 h after HRG; Fig. 2A, B). Additionally, gene set enrichment analysis (GSEA) using normalized enrichment scores (*49*) did not show any significant enrichment of c-Myc-target genes during 12–16 h (Fig. 4C). However, CDK4i induced a significant enrichment of c-Myc-target genes at 16 h after HRG stimulation, followed by a downregulation at 24–28 h compared to that in the control (Fig. 4D). We analyzed the temporal expression of cell cycle-related c-Myc-target genes.

Focusing on genes exhibiting a higher abundance of H3K27Ac modifications in their promoter than their enhancer region (Fig. S4D). *CCNE1* expression was significantly enhanced 16 h after HRG stimulation and CDK4i treatment, whereas the expression of *CCND1*, *CDK1*, *CDC25A*, *E2F2*, *E2F3*, and *CDKN1B* showed delayed upregulation during 20–28 h. These findings suggested that CDK4 inhibition alters the H3K27Ac levels in the promoter regions of c-Myc-target genes near the R point, promoting the expression of cell cycle-related genes. Therefore, when the G1/S transition, originally regulated by the cyclin D1 and CDK4 complex, is inhibited by CDK4i, it is maintained by an alternative pathway facilitated by c-Myc activity.

We confirmed whether the delayed G1/S entry following CDK4i treatment was due to c-Myc activation by testing the effects of an inhibitor of c-Myc transcriptional activity (EN4). In both MCF-7 and T47D cells after 30 h, the combination of CDK4i and EN4 increased the proportion of G1-phase cells and decreased the number of S/G2/M-phase cells in a dose-dependent manner (Fig. 4E–H), while no change was observed with EN4 alone (Fig. S4E–H). These results suggested that the transcriptional activity of c-Myc promotes the delayed G1/S entry and escape from cell cycle arrest in response to CDK4 inhibition.

To further validate these findings using clinical data, we retrospectively analyzed the microarray data from patients with luminal A subtype breast cancer who received long-term (16 weeks, *n* = 23) or short-term (4 weeks, *n* = 26) CDK4/6i therapy (*50*). *PCNA*, a target gene for both c-Myc and E2F involved in DNA replication during G1/S transition, showed a 0.5 log2 fold change (FC) between the long-term and short-term treatment groups (Fig. 4I). *PCNA* mRNA expression increased after CDK4i treatment from 4 to 12 h (16–24 h after HRG) relative to the control (Fig. 4J). Collectively, these findings suggested that CDK4 inhibition-induced epigenetic modifications activated c-Myc-target genes, ultimately promoting cell cycle maintenance in cancer cells (Fig. 4K).

### CDK4i induces ErbB2 degradation and delays G1/S transition

c-Myc is known to promote G1/S transition and cell proliferation (*25*, *35*). Therefore, if the transcriptional activity of c-Myc is enhanced by CDK4i at 16 h, the time to G1/S transition is unlikely to be extended; however, the G1/S transition was prolonged in our study (Fig. 3D, E; Fig. 4A). We found that the levels of phosphorylated ErbB2, ERK, and AKT, which induce *c-Myc* mRNA expression and stabilize the c-Myc protein, decreased after CDK4 inhibition, leading to a slight reduction in c-Myc protein level (Fig. 5A, B). Although the expression of ErbB2 was lower in MCF-7 and T47D cells than in HER2-positive cells, growth in these cell lines is mainly dependent on ErbB2-ErbB3 heterodimer activation after HRG stimulation given their low expression of EGFR and ErbB4 (*37*). This finding suggested that the CDK4i-induced downregulation of the ErbB2 pathway may prolong the G1/S transition in MCF-7 and T47D cells (Fig. 3D; Fig. S3F). To investigate this mechanism, we focused on the Hsp90-chaperone system, particularly with respect to the stability and maturation of nascent ErbB2. As WT MCF-7 cells naturally express low levels of ErbB2, we constructed MCF-7 cells with moderate overexpression of ErbB2 (Fig. S5A, B). After confirming the unaltered protein expression of ErbB2, PY100 (pan-tyrosine phosphorylation for estimating p-ErbB2 level), and Hsp90 under CDK4i (Fig. S5C–G), we examined the co-localization patterns using Mander’s and Pearson’s coefficients. Compared to control conditions, CDK4i reduced the co-localization of Hsp90 with ErbB2 or PY100 (Fig. 5C–F). No significant difference was observed in the total number of nuclei (Fig. S5H, I). These results suggested that CDK4i inhibits the interaction between Hsp90 and ErbB2, leading to ErbB2 misfolding. In addition, ErbB2 protein expression was reduced 20 h after HRG stimulation (Fig. 5G, H; Fig. S5J); however, *ErbB2* mRNA levels did not change regardless of CDK4i treatment (Fig. S5K).

**Fig. 5.**
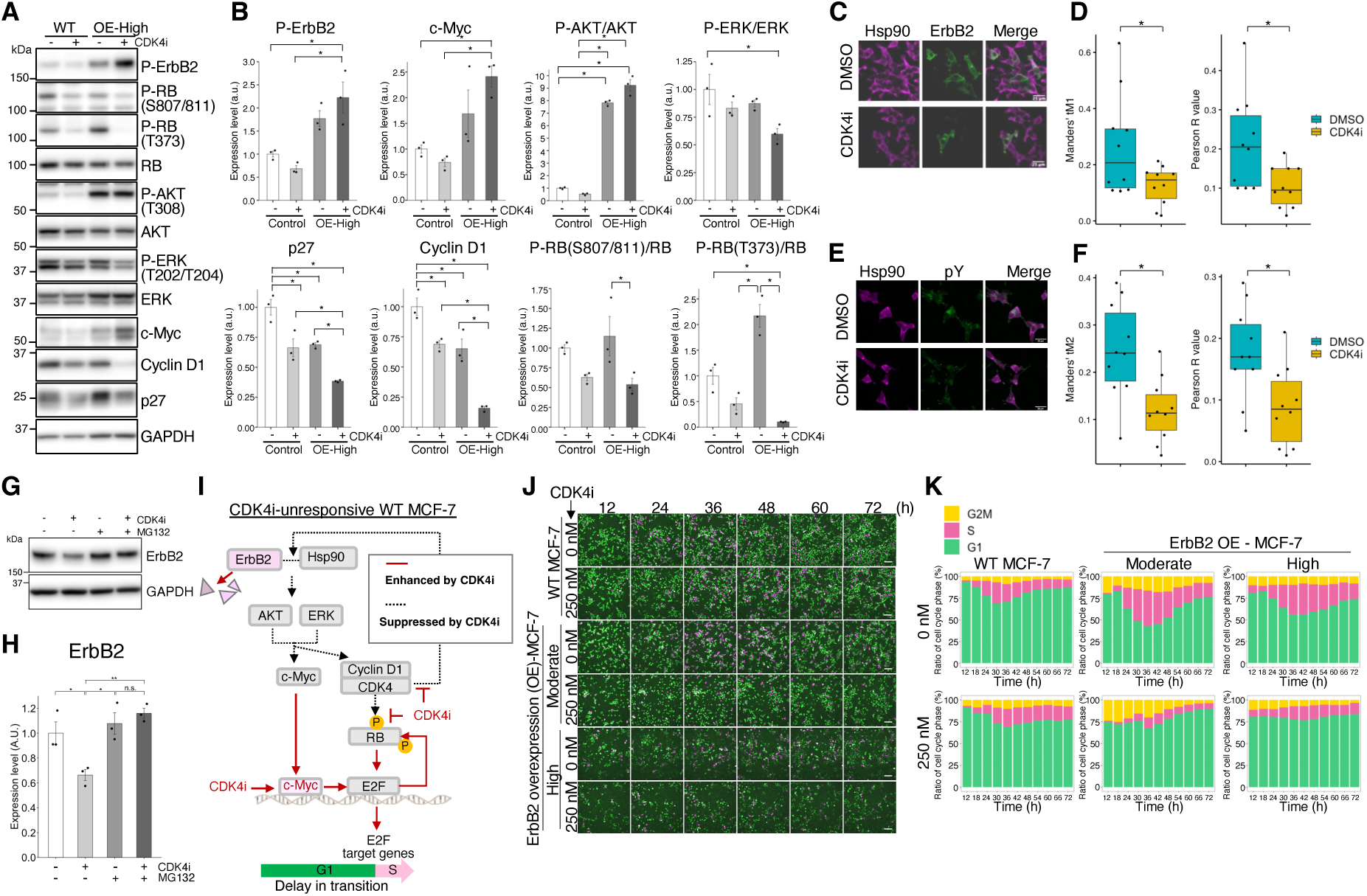
High ErbB2 expression induces cell cycle arrest by repressing cyclin D1 expression via c-Myc under CDK4 inhibition. (**A, B**) WT MCF-7 and ErbB2-overexpressing MCF-7 cells (high levels) were treated with CDK4i or DMSO 12 h after HRG stimulation and harvested 24 h after HRG stimulation. Western blotting of (A) p-ErbB2, p-RB (Ser^807/811^), p-RB (Thr^373^), RB, p-AKT (Thr^308^), AKT, p-ERK (Thr^202^/Thr^204^), ERK, c-Myc, cyclin D1, and p27 expression (in control cells, i.e., DMSO treatment) after normalization with GAPDH expression (B). (**C–F**) ErbB2-overexpressing cells (moderate) were individually treated with 250 nM CDK4i or DMSO (no treatment) 12 h after stimulation with 10 nM HRG and fixed 16 h after stimulation with 10 nM HRG (CDK4i or DMSO addition 4 h later). Hsp90 and ErbB2 were co-immunostained. Scale bar: 25 µm (C). Co-localization rate of Hsp90 and ErbB2 based on Mander’s tM1 and Pearson’s R values determined using Coloc 2 plugin of Image J; *n* = 10 (D). Hsp90 and PY100 were co-immunostained. Scale bar: 25 µm (E). Co-localization rate of Hsp90 and PY100 based on Mander’s tM2 and Pearson’s R values; *n* = 10 (F). (**G, H**) FUCCI stably expressing MCF-7 cells were treated with CDK4i or DMSO (control) as well as 10 mM MG132 12 h after stimulation with 10 nM HRG and collected 8 h later. (**G**) Expression of ErbB2 and GAPDH determined by western blotting. (**H**) ErbB2 expression normalized to GAPDH expression; *n* = 3. (**I**) In CDK4i-unresponsive WT MCF-7 cells, the loss of ErbB2 protein stability impairs the regulation of c-Myc-mediated G1/S transition, leading to the delayed G1/S transition. (**J**) WT MCF-7 and ErbB2-overexpressing cells (moderate and high levels) were individually treated with 250 nM CDK4i or DMSO (no treatment) 12 h after stimulation with 10 nM HRG, and images were acquired every 20 min until 72 h after stimulation. Scale bar: 100 µm. (**K**) Images in (J) were analyzed to determine the proportion (%) of each cell cycle phase in the entire set of images; *n* = 3. In (B) and (H), \**P* < 0.05 (Tukey’s test). In (D) and (F), \**P* < 0.05 (with respect to control and CDK4i; *t*-test).

Therefore, we investigated whether the downregulation of ErbB2 was caused by its protein degradation using the proteasome inhibitor MG132. Upon MG132 treatment, ErbB2 protein expression remained unchanged (Fig. 5G, H). This observation indicated that CDK4i disrupts the interaction between Hsp90 and ErbB2, triggering ErbB2 degradation (Fig. S5L). To confirm this, we treated cells with the Hsp90 inhibitor geldanamycin at a concentration that suppressed ErbB2 expression by 40%, similar to the effect of CDK4i (Fig. S5M, N). As a result, cell cycle progression grounded to a halt, and there was no substantial delay in the G1/S transition (Movie S1). Therefore, the loss of ErbB2 protein stability following CDK4i treatment may impair the regulation of c-Myc-mediated G1/S transition, even if c-Myc is activated by epigenetic modification, thereby delaying the G1/S transition (Fig. 5I).

### High ErbB2 expression induces cell cycle arrest by repressing cyclin D1 activity via c-Myc under CDK4 inhibition

We speculated that ErbB2 protein instability contributes to the delay in G1/S entry under CDK4i and thus could be counteracted by ErbB2 overexpression. To test this hypothesis, we generated MCF-7 cells expressing varying levels of ErbB2, from moderate to high (Fig. S5A, B), and analyzed their cell cycle phase distribution using live-cell imaging (Fig. 5J). After CDK4i treatment, the proportion of S/G2/M-phase cells with high ErbB2 expression was significantly reduced (OE-Moderate: 14.9%, OE-High: 12.5%; Fig. 5K), indicating cell cycle arrest. We examined the protein expression of cyclin D1 and p-RB, which are critical for initiating cell cycle entry (Fig. 5A, B). We observed a significant reduction in both cyclin D1 and p-RB levels. Therefore, we investigated the cause of the decreased cyclin D1 levels. The p27 protein reportedly promotes the assembly of the cyclin D1/CDK4 complex in the G1 phase (*34*), while its gene expression is inhibited by p-AKT and c-Myc (*35*). Our data revealed that p27 protein expression decreased after CDK4i treatment, whereas p-AKT and c-Myc protein levels tended to increase post treatment. By contrast, p-ERK levels remained unchanged under different conditions. These findings suggested that, in ErbB2-overexpressing cells, the enhanced p-AKT–c-Myc axis antagonizes the protein expression of p27 and cyclin D1, leading to cell cycle arrest in the absence of CDK4 activity.

### Reversible cell cycle arrest is maintained by c-Myc transcriptional activity

Drug resistance is closely linked to a reversible cell cycle arrest state regulated by residual survival signals (*11*). In ErbB2-overexpressing cells, a cell cycle arrest state is induced after CDK4 inhibition, but this may be reversed by the survival signals from c-Myc. In CDK4i-treated cells with high ErbB2 expression, CDK4i removal increased cyclin D1, p27, and p-RB levels, while c-Myc expression remained unchanged (Fig. 6A, B). We analyzed the c-Myc phosphorylation (Ser^62^) levels to confirm its elevated transcriptional activity in cell cycle arrested cells. Single-cell analysis revealed that CDK4i depletion decreased the ratio of phosphorylated c-Myc (Ser^62^) to cyclin D1 (p-c-Myc/cyclin D1) and increased the ratio of S-phase cells from 18.7% to 47.5% (Fig. 6C). ErbB2-overexpressing cells maintained a higher p-c-Myc/cyclin D1 ratio than wild-type cells (Fig. S6A), suggesting that sustained cell cycle arrest may result from persistent c-Myc transcriptional activation.

**Fig. 6.**
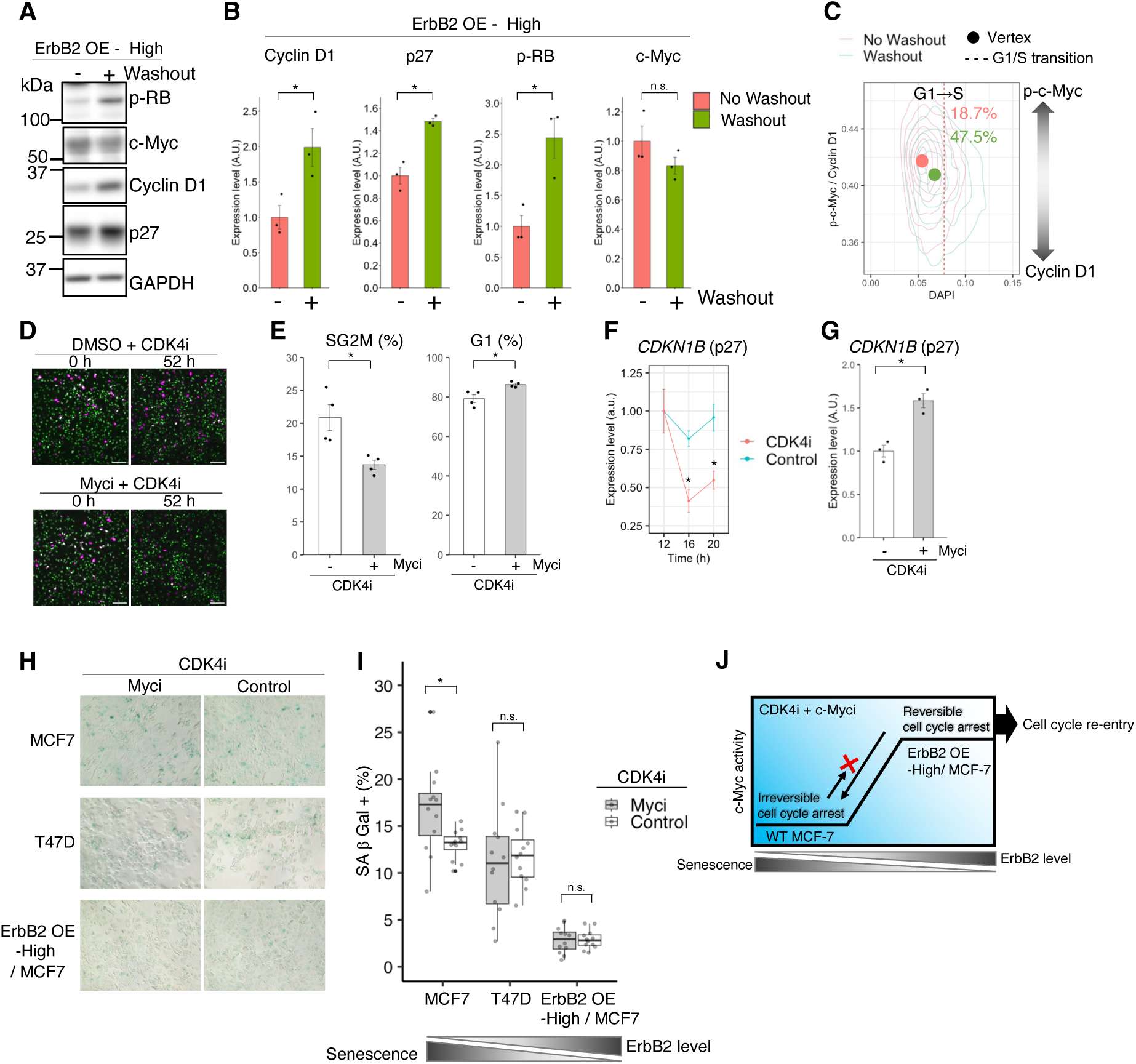
Reversible cell cycle arrest is maintained under c-Myc transcriptional activity. (**A–G**) MCF-7 cells overexpressing ErbB2 (high levels). (**A, B**) Cells were treated with 250 nM CDK4i 12 h after HRG stimulation, subjected to a wash/no-wash treatment 4 h later, and harvested 40 h after HRG stimulation. Western blotting results showing the expression of p-RB, c-Myc, cyclin D1, p27, and GAPDH. After individually normalizing the proteins relative to GAPDH, the ratio to the no-wash condition was quantified; *n* = 3. (**C**) Washing operations were performed under the same conditions as in (A, B); cells were fixed 40 h after HRG stimulation and immunostained with p-c-Myc and cyclin D1 antibodies together with DAPI; *n* = 3 859. A higher value on the y-axis indicates that the amount of p-c-Myc in one cell was greater than that of cyclin D1. The circle shows the vertex of each histogram, and the red dotted line indicates the G1/S transition time point. (**D, E**) Cells were pre-treated for 30 min with/without 64 nM c-Myc inhibitor (EN4) 11.5 h after HRG stimulation. The cells were then incubated with 250 nM CDK4i for 8 h, washed, and imaged 52 h later (72 h after HRG stimulation, D). (**E**) Percentage distributions of G1 and S/G2/M phases based on the images in (D); *n* = 3. Scale bar: 100 µm. (**F**) Cells were treated with 250 nM CDK4i or DMSO (control) 12 h after HRG stimulation and p27 mRNA levels up to 20 h after HRG stimulation were examined by qPCR; *n* = 3. (**G**) After the cells were treated with or without EN4 under the same conditions as in (D–E), RNA was collected 8 h after treatment with inhibitors, and *CDKN1B* mRNA levels were examined by qPCR; *n* = 3. (**H**) WT MCF-7, WT T47D, and ErbB2 (high) OE MCF-7 cells were pre-treated for 30 min with/without 4 mM c-Myc inhibitor (EN4) 11.5 h after HRG stimulation, incubated with 250 nM CDK4i for 6 d, and SA-β-Gal staining was assessed. (**I**) Percentage SA-β-Gal-stained cells based on the images in (H); *n* = 12. (**J**) ErbB2 levels modulate the irreversibility of the G1/S transition under CDK4i treatment. In (B), (E), (F), (G), (I) \**P* < 0.05 (with respect to control and each inhibitor; *t*-test).

To investigate whether blocking c-Myc transcriptional activity could prevent reversible cell cycle arrest, we treated cells with EN4 and CDK4i for 8 h, washed the inhibitors, and analyzed cell cycle distribution at 52 h (72 h after HRG stimulation; Fig. 6D, E). The combined treatment increased G1-phase and reduced S/G2/M-phase cell numbers, indicating impaired cell cycle re-entry. CDK4i reduced p27 (*CDKN1B*) mRNA levels, which were restored by co-treatment with EN4 (Fig. 6F, G), suggesting direct c-Myc regulation of p27 gene expression. Overall, these data supported the hypothesis that, in cells overexpressing ErbB2, CDK4 inhibition promotes reversible cell cycle arrest due to an excessive increase in c-Myc transcriptional activity and rapid decrease in cyclin D1 protein expression, which is attributed to the reduction in p27 gene and protein expression.

Given the association between stable cell cycle arrest and cellular (*51*, *52*) or ErbB2-induced senescence (*53*), we examined the senescence phenotype. Senescence associated β-galactosidase (SA-β-Gal) staining intensity after 6 d of CDK4i treatment showed an inverse correlation with ErbB2 expression. EN4 (4 µM) enhanced SA-β-Gal staining in WT MCF-7 cells beyond the effect of CDK4i alone but showed no additional effect in T47D or high ErbB2-expressing MCF-7 cells (Fig. 6H, I). This finding was consistent with cell line-specific sensitivity, since 2 nM EN4 induced cell cycle arrest in WT MCF-7 cells, whereas 2 µM was required for WT T47D cells (Fig. 4E, F). Our results suggested that ErbB2 levels modulate the irreversibility of the G1/S transition under CDK4i treatment, and the addition of a c-Myc inhibitor supports the irreversible state (Fig. 6J).

## Discussion

Our findings reveal that ErbB2 receptor levels modulate the relative roles of cyclin D1 and c-Myc in the G1/S transition, ultimately providing two distinct pathways for cell cycle progression or arrest in the presence of CDK4i.

The luminal A subtype accounts for 30% of breast cancer cases. Although these patients typically receive standard hormone therapy, CDK4/6is are also commonly used in cases of more advanced or metastatic tumors (*5*, *54*). However, their success is limited by heterogeneous drug responses. Notably, an ErbB2-overexpressing subpopulation has been identified within luminal A breast cancer cells, which were initially characterized by low ErbB2 expression (*55*). Given that these subpopulations occur relatively infrequently, at approximately 7% of the total population (*56*), identifying differences in cell cycle regulation, especially in terms of ErbB2 expression, is critical for improving cancer treatment. Long-term CDK4i treatment, commonly used for luminal A breast cancer, activates alternative pathways, such as PI3K/AKT/mTOR, androgen receptor, and Hippo signaling (*13*, *14*, *16*, *17*), along with transcriptional reprogramming via non-coding RNAs (*15*, *57*), promoting the development of drug resistance. To mitigate this resistance, pathways that may act at the very early step of cell cycle entry must be inhibited. Accordingly, we investigated how ErbB2 levels within cells of the same luminal A subtype affect the core cell cycle regulatory network and CDK4 inhibitor sensitivity (Fig. 7A, B). In MCF-7 cells under HRG stimulation, CDK4 RNA levels increased and remained elevated through R-point and S-phase entry, whereas CDK6 RNA levels were quite low compared to those of CDK4 and tended to decrease after HRG stimulation (Fig. S4D). Therefore, we focused on the effect of CDK4 inhibitors rather than CDK4/6 inhibitors.

**Fig. 7.**
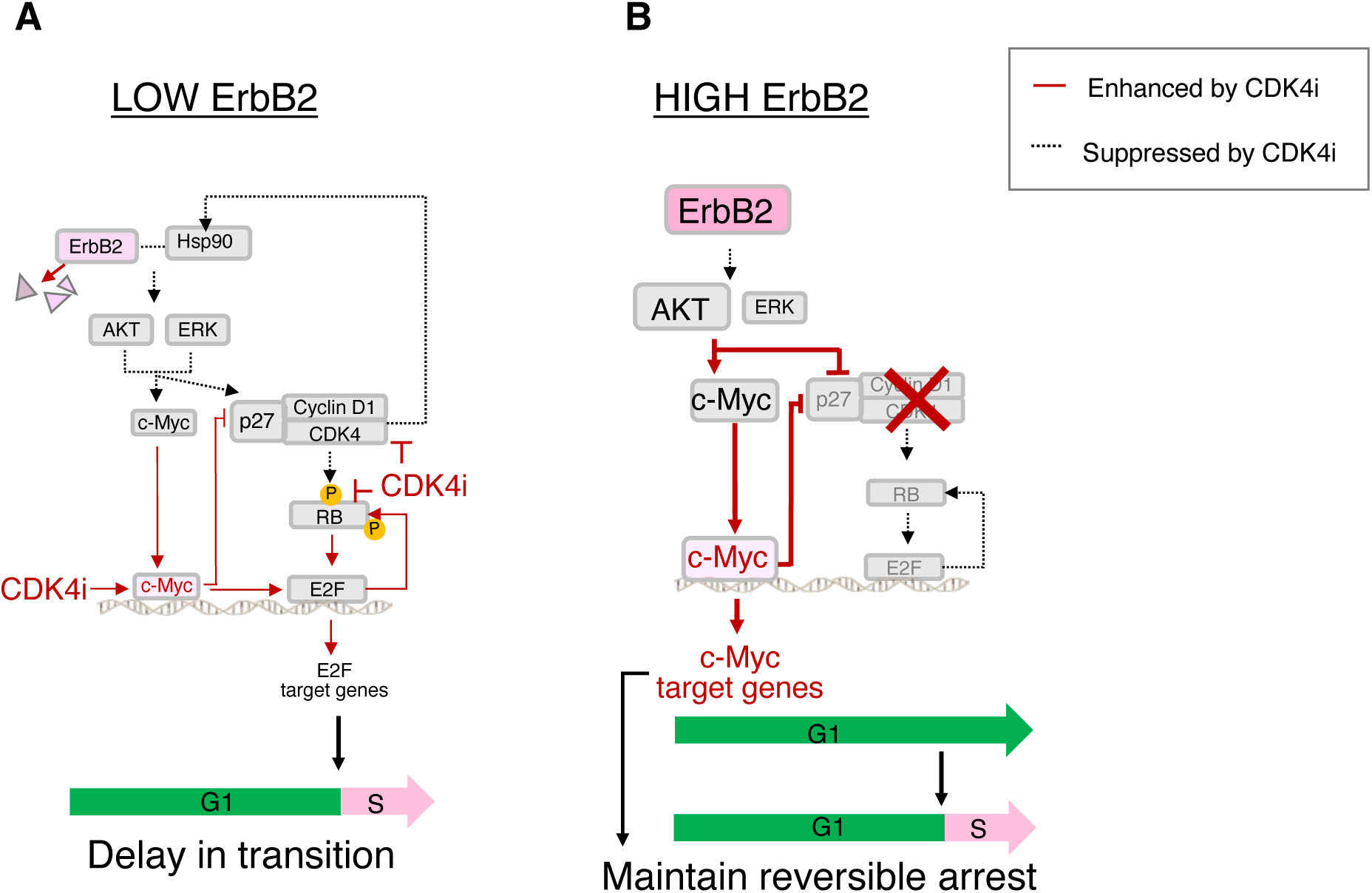
Proposed model the ErbB2 receptor regulation during G1/S transition via cyclin D1 and c-Myc in CDK4i non-responsive cells. (**A**) In low ErbB2-expressing cells, CDK4 inhibition leads to reduced pathway activity due to ErbB2 degradation via Hsp90 discolocalization. This attenuates the transcriptional activity of c-Myc activated by epigenetic changes. Consequently, a subpopulation of cells unresponsive to CDK4i exhibits a delayed G1/S transition. (**B**) In high ErbB2-expressing cells, increased AKT activity leads to increased c-Myc expression. Subsequently, the expression of p27, which is critical for the stability of cyclin D1/CDK4 during the G1 phase, is suppressed by c-Myc, resulting in cell cycle arrest after CDK4 inhibition. This arrest is reversibly maintained by the strong transcriptional activity of c-Myc, ultimately sustaining survival signals.

The temporal dynamics of the cell cycle regulators and live-cell imaging analysis showed that the roles of cyclin D1 and c-Myc, key regulators of the G1/S transition, vary depending on the expression status of ErbB2 under CDK4i treatment. This variation ultimately affects the timing and irreversibility of the G1/S transition. In MCF-7 cells, a representative luminal A breast cancer cell line with low ErbB2 expression, the G1/S transition is normally regulated by the cyclin D1/CDK4 complex via Hsp90 (Fig. 5C–E; Fig. S5C–O). Although CDK inhibitors are known to sequester Hsp90 and CDK4/6 (*58*), our results suggested that the function of Hsp90 is complemented by CDK4 activity and contributes to stabilizing the ErbB2 protein. This positive feedback mechanism provides valuable insights into the signal-dependent complexity of cell cycle regulation. In the non-responsive subpopulations to CDK4 inhibition, the G1/S transition persists, but with delays due to epigenetic modification and increased c-Myc transcriptional activity and ErbB2 degradation (Fig. 4; Fig. 7A).

However, the excessive transcriptional activity of c-Myc promotes cell cycle arrest (Fig. 6A–G) by rapidly reducing cyclin D1 and p27 levels. The expression of p27, which recruits cyclin D1 to the nucleus during the G1 phase, is also repressed by AKT and c-Myc (*35*). Therefore, in high ErbB2-expressing MCF-7 cells, with high levels of AKT and c-Myc activity, CDK4 inhibition increases free cyclin D1 levels, rendering it more susceptible to degradation (*59–61*). This process generates a hypophosphorylated RB state in ErbB2-overexpressing MCF-7 cells compared to WT cells, leading to cell cycle arrest due to a decrease in free E2F now bound to c-Myc (Fig. 7B).

c-Myc regulates the cell survival pathway (*35*, *62*). In ErbB2-overexpressing MCF-7 cells, cell cycle arrest was reversible after CDK4 inhibition (Fig. 6D, E), suggesting that the transcriptional activity of c-Myc promotes sustained cell survival in this context. Co-treatment with inhibitors targeting the transcriptional activity of c-Myc and CDK4 may be necessary to induce irreversible cell cycle arrest (Fig. 6H, I). However, at the same concentration at which the Myc inhibitor promoted cellular senescence in WT MCF-7 cells, the percentage of senescent WT T47D and ErbB2-overexpressing cells did not change compared to that in the respective control conditions. For drug response assessments, the threshold at which c-Myc switches from supporting cell survival to inducing irreversible cell cycle arrest in relation to ErbB2 levels could inform the development of more effective molecular-targeted treatment strategies for breast cancer.

Overall, our results suggest that the c-Myc-to-cyclin D1 ratio provides valuable quantitative insights into the varying sensitivity of cancer cells to CDK4is, without relying solely on observable cell proliferation rates or single biomarker gene analysis.

## Materials and methods

### Reagents

The MEKi PD0325901 was purchased from FUJIFILM Wako Pure Chemical Corp. (Osaka, Japan); CDK4i V was purchased from Calbiochem (San Diego, CA, USA); AKTi VIII was purchased from Cayman Chemical (Ann Arbor, MI, USA); EN4 (Myci) was purchased from Selleck Chemicals (Houston, TX, USA); geldanamycin was purchased from FUJIFILM Wako Pure Chemical Corp.; blasticidin was purchased from Kaken Pharmaceutical (Tokyo, Japan); Lipofectamine LTX was purchased from Thermo Fisher Scientific (Waltham, MA, USA); and HRG was procured from R&D Systems (Minneapolis, MN, USA).

### ErbB2 overexpression

MCF-7 (ATCC, HTB-22) cells overexpressing full-length human ErbB2 (HER2) were established as follows: Human ErbB2 cDNA in the pCMV6-XL5 vector was purchased from OriGene (Catalog No: TC128161, Rockville, MD, USA). The cDNA fragment was cloned into the vector pCMV-6-Neo vector (OriGene). The plasmid was then transfected into MCF-7 cells using Lipofectamine LTX (Invitrogen, Carlsbad, CA, USA) according to the manufacturer’s protocol, and stable transfectants were selected with G418 (neomycin). Colonies derived from single cells compartmentalized in 96-well plates were isolated and examined. ErbB2 expression was examined via western blotting to group cells with moderate or high ErbB2 expression (as described below for protein extraction and immunoblotting).

### Cell culture

MCF-7, ErbB2-overexpressing MCF-7, and T47D (ATCC, HTB-133) cells were cultured in Dulbecco’s modified Eagle’s medium (DMEM, Nacalai Tesque, Kyoto, Japan) supplemented with 10% (*v/v*) fetal bovine serum (FBS, Sigma-Aldrich, Saint Louis, MO, USA) and 1% penicillin–streptomycin (100 IU/ml penicillin, 100 μg/ml streptomycin; Nacalai Tesque) at 37 °C in a humidified 5%-CO2 atmosphere. Cells were routinely tested using a CycleavePCR Mycoplasma Detection Kit (Takara bio, CY-232); no mycoplasma contamination was detected.

### FUCCI transfection

MCF-7 and ErbB2-overexpressing MCF-7 cells stably expressing the FUCCI biosensor were generated using a PiggyBac transposase system (*63*). Lipofectamine LTX (Thermo Fisher Scientific) was used to transfect the FUCCI plasmid (pPBbsr2-H2B-iRFP-P2A-mScarlet-I-hGem-P2A-PIP-tag-NLS-mNeonGreen; https://benchling.com/s/seq-LPZ1tLdpgnpJIYOG2ujR) along with hyPBase (https://benchling.com/s/seq-oGkw53b41IZqvzF5yQ9K) into MCF-7 and ErbB2-overexpressing MCF-7 cells, following the manufacturer’s protocol (*64*). Cells were suspended in a drug-free medium for 48 h and treated with blasticidin (10 µg/ml) to separate those stably expressing the plasmid. Colonies derived from single cells compartmentalized in 96-well plates were selected by microscopic live-cell observation of Geminin and PIP-tag fluorescence.

### Protein extraction and immunoblotting

MCF-7, T47D, and ErbB2-overexpressing (moderate or high) MCF-7 cells were seeded in six-well plates (450 000 cells/well) and cultured in 10% FBS containing DMEM for 48 h. The medium was changed to FBS-free phenol red-free DMEM (FUJIFILM Wako Pure Chemical Corp.) containing 1% penicillin–streptomycin (100 IU/ml penicillin, 100 μg/ml streptomycin; Nacalai Tesque) and 1% 100 mmol/L sodium pyruvate solution (FUJIFILM Wako Pure Chemical Corp.), and the cells were cultured for 17 h. The medium was changed again, and the cells were cultured for 3 h. After administration of 10 nM HRG (R&D Systems) for 12 h, the cells were independently treated with CDK4i and dimethyl sulfoxide (DMSO). Serum was not included under conditions where HRG was added. Total cell lysates were prepared in RIPA buffer (Nacalai Tesque, 1648834) containing protease inhibitors (11836153001, Roche Rotkreuz, Switzerland) and phosphatase inhibitors (Roche, 4906837001) at 4 °C. Cells were vortexed for 5 min and centrifuged at 16 000 × *g* and 4 °C for 15 min. The supernatant was collected and denatured by heating at 95 °C with sample buffer (5×) and 2-mercaptoethanol (Nacalai Tesque, 21418-55) for 5 min. The proteins were subjected to SDS polyacrylamide gel electrophoresis (Nacalai Tesque, 1306644), transferred to polyvinylidene fluoride membranes (Merck, Darmstadt, Germany), and incubated with 10% blocking buffer (Nacalai Tesque, 03953-95) prepared in Tris-buffered saline/Tween 20 (TBS-T; T9142; Takara Bio, Shiga, Japan) for 1 h at 25 °C. Membranes were probed overnight with antibodies against AKT (1:1 000; #4685; Cell Signaling Technology, Danvers, MA, USA), p-AKT (Thr^308^; 1:1 000; Cell Signaling Technology; #2965), ERK (1:1 000; Cell Signaling Technology; #9102), p-ERK (Thr^202^ Tyr^204^; 1:1 000; Cell Signaling Technology; #4370), HER2/ErbB2 (1:1000; Cell Signaling Technology; #2165), p-HER2/ErbB2 (Thr^1221/1222^; 1:1 000; Cell Signaling Technology; #2249), p-c-Fos (Ser^374^; 1:1 000; #ab55836; Abcam, Cambridge, UK), c-Fos (1:200; #sc-166940; Santa Cruz Biotechnology, Dallas, TX, USA), p-c-Myc (Ser^62^; 1:1 000; Abcam; #ab51156), c-Myc (1:200; Santa Cruz; #sc-40), E2F-1 (1:1 000; Cell Signaling Technology; #3742), p-RB (Ser^807/811^; 1:1 000; Cell Signaling Technology; #8516), p-RB (Thr^373^; 1:1 000; Abcam; #ab52975), RB1 (1:1000; #SPM353; NOVUS biologicals, Centennial, CO, USA), cyclin D1 (1:1000; Cell Signaling Technology; #2978), cyclin E (1:200; Santa Cruz; #sc-247), cyclin A (1:200; Santa Cruz; #sc-271682), CDK4 (1:1 000; Santa Cruz; #DSC-35), CDK2 (1:1 000; Santa Cruz; #D-12), P27 Kip1 (1:1 000; Cell Signaling Technology; #3686), and GAPDH (1:4 000; 10494-1-AP; Proteintech, Boston, MA, USA) in TBS-T including 10% blocking buffer at 4 °C. They were then incubated with horseradish peroxidase (HRP)-conjugated secondary antibodies, HRP-conjugated anti-mouse IgG (1:5000; Cell Signaling Technology; #7076) and HRP-conjugated anti-rabbit IgG (1:5000; Cell Signaling Technology; #7074), for 1 h at room temperature. The membranes were developed using a chemiluminescent HRP substrate (Merck). Protein bands were detected using an Amersham Imager 600 UV (GE Healthcare, Eindhoven, Netherlands) and visualized using the ImageJ-Fiji software (v.2.14.0/1.54f, NIH, Bethesda, MD, USA).

### qRT-PCR

qRT-PCR was conducted as previously described (*65*). RNA was isolated from cells using NucleoSpin RNA Plus (Cat. No. 740984.250; Macherey-Nagel, Duren, Germany) according to the manufacturer’s protocol. cDNA was synthesized using ReverTra Ace qPCR RT Master Mix (TOYOBO, Osaka, Japan), and quantitative PCR was performed using a KOD SYBR qPCR kit (TOYOBO) on a CFX96 Real-Time PCR System (Bio-Rad Laboratories, Hercules, CA, USA). The ΔΔCq method was used to quantify gene expression, using *RPL27* expression as an internal reference (*66*). The primers used for real-time PCR are listed in Table S1.

### Cell cycle analysis using the FUCCI biosensor

Cell cycle analysis was conducted as previously described (*65*). Time-lapse microscopy was performed using an IN Cell Analyzer 2500HS (GE Healthcare, Chicago, IL, USA) equipped with a CFI S Plan Fluor ELWD dry objective lens 20× (NA: 0.45) at an excitation wavelength of 475 nm and emission wavelength of 511 nm for PIP-tag-mNeonGreen, and an excitation wavelength of 542 nm and emission wavelength of 597 nm for mScarlet-hGem. Cells were maintained at 37 °C in a humidified 5%-CO2 atmosphere and imaged at 20-min intervals for 72 h. Image processing and quantification of cell cycle phase distribution were performed using ImageJ-Fiji software (v.2.14.0/1.54f, NIH). Masks of the mScarlet-hGem and PIP-tag-mNeonGreen signals were obtained by applying a fixed threshold using “Analyze Particles” and automatically counted as ROIs. Cell cycle phases were determined in the following manner: G1 phase, PIP-tag-positive; S phase, hGem-positive; G2/M phase, PIP-tag- and hGem-positive. The average brightness of mScarlet and mNeonGreen was obtained using cell-tracking software (Olympus, Tokyo, Japan). The threshold parameters for mScarlet and mNeonGreen were set manually for each cell line. We employed an algorithm for estimating the cell cycle phase at the single-cell level using R packages [brunnermunzel (v.1.4.1), multimode (v.1.4), rlist (v.0.4.6.1), and tidyverse (v.1.3.0)]. The duration of each cell cycle phase was obtained from the single-cell data.

### Nucleus count

The number of nuclei was quantified based on the fluorescence of the FUCCI probes, and signals for a nuclear marker (histone H2B), mScarlet-hGem, and PIP-tag-mNeonGreen were used to determine cell proliferation. Image processing and nuclear ratio quantification were performed using the ImageJ-Fiji software (v.2.14.0/1.54f, NIH). Masks of the merged nuclear marker, mScarlet-hGem, and PIP-tag-mNeonGreen signals were obtained by applying a fixed threshold using “Analyze Particles” and automatically counted as ROIs. The nuclear ratio was calculated by dividing the number of nuclei at the end of observation by that at the beginning. Thus, a nuclear ratio near 1.0 indicated potential cell cycle arrest, above 1.0 indicated cell proliferation, and below 1.0 indicated apoptosis.

### Immunostaining and quantitative analysis of images

MCF-7 and ErbB2-overexpressing (moderate or high) cells were seeded in 12-well plates (200 000 cells/well) and cultured in 10% FBS containing DMEM for 48 h. The medium was changed to FBS-free phenol red-free DMEM (FUJIFILM Wako Pure Chemical Corp.) containing 1% penicillin–streptomycin (100 IU/ml penicillin, 100 μg/ml streptomycin; Nacalai Tesque) and 1% 100 mmol/L sodium pyruvate solution (FUJIFILM Wako Pure Chemical Corp.), and the cells were cultured for 17 h. The medium was changed again, and the cells were cultured for 3 h. After administration of 10 nM HRG (R&D Systems) for 12 h, the cells were independently treated with CDK4i and DMSO. Serum was not included under conditions where HRG was added. Cells were washed twice with PBS at the indicated times, fixed for 2 min with pre-prepared methanol at –20 °C, washed twice with Blocking One (Nacalai Tesque, 03953-95), and incubated overnight at 4 °C with Blocking One. The next day, cells were incubated overnight at 4 °C with primary antibodies diluted in 10% goat serum (Thermo Fisher Scientific, 16210064)/Blocking One, washed five times with PBS, and incubated with fluorescent dye-conjugated secondary antibody, 2 μg/ml DAPI (Nacalai Tesque, 11034-56), and 0.5 μg/ml CellMask (Thermo Fisher Scientific, H32721) for 1 h at room temperature. Cells were washed five times with PBS, and images were acquired using the IN Cell Analyzer 2500HS (GE Healthcare Life Science) at 20× magnification. At least three images were collected for each treatment, containing at least 100 cells each, for quantification using the ImageJ-Fiji software (v.2.14.0/1.54f, NIH) and CellProfiler (v.4.2.1). Co-localization analysis for Hsp90 with either ErbB2 or PY100 was performed using the “Coloc 2” plugin in the ImageJ-Fiji software. Cell fractionation was performed on CellMask-stained images using CellProfiler, and the mean intensity was calculated for p-c-Myc, cyclin D1, and DAPI in each cell. The ratio of phosphorylated c-Myc (Ser^62^) to cyclin D1 (p-c-Myc/cyclin D1) for each cell was plotted over the DAPI intensity. The DAPI intensity indicating the G1/S transition was determined based on the percentage of cells that were in the S/G2 /M phase 24 h after HRG stimulation. In WT MCF-7 cells, 22.4% of cells were in the S/G2/M phase, at which time the DAPI intensity was 0.105. In MCF-7 cells overexpressing ErbB2 (high), 25.4% of the cells were in the S/G2/M phase, at which time the DAPI intensity was 0.0775.

### Analysis of senescence associated β-galactosidase-positive cells

SA-b-gal staining was conducted as previously described (*65*). Briefly, senescent cells were stained using a Senescence β-Galactosidase Staining Kit (#9860; Cell Signaling Technology) according to the manufacturer’s instructions. Images were acquired with a BZ-9000 microscope (Keyence, Osaka, Japan) at 20× magnification. The percentage of SA-β-gal-positive cells was calculated from images using CellProfiler (v.4.2.1). The total number of cells was determined by visualizing nuclear segmentation with DAPI staining. The number of SA-β-gal-stained cells was identified using the UnmixColors module. The percentage of SA-β-gal-positive cells was calculated by dividing the number of SA-β-gal-stained cells by the total number of nuclei in each field of view.

### RNA-seq

FUCCI-expressing MCF-7 cells were seeded in six-well plates (450 000 cells/well) and cultured in 10% FBS containing DMEM for 48 h. The medium was changed to FBS-free phenol red-free DMEM, the cells were cultured for 17 h, the medium was changed again, and the cells were cultured for 3 h. After administration of 10 nM HRG for 12 h, the cells were independently treated with CDK4i and DMSO. Serum was not included under conditions where HRG was added. Cells were washed twice with PBS at the indicated times. Total RNA was isolated using NucleoSpin RNA Plus (Macherey-Nagel; Cat. No. 740984.250) at the specified time. Library construction and RNA-seq were performed on an Illumina sequencing platform (Rhelixa Tokyo, Japan). Poly(A) RNA was prepared with a Poly(A) mRNA Magnetic Isolation Module (E7490; New England Biolabs, Ipswich, MA, USA), and the sequencing library was generated using a NEBNext Ultra II Directional RNA Library Prep Kit (New England Biolabs, E7760). RNA-seq was performed on a NovaSeq 6000 (Illumina, San Diego, CA, USA) with 150-bp paired-end reads. Raw reads were quality-checked using FastQC (v.0.11.9) and trimmed using TrimGalore (v.0.6.0) (*67*). Trimmed reads were mapped using STAR (v.2.7.9a) (*68*) with GRCh38 as the reference genome. featureCounts (v.1.6.4) (*69*) was used to construct the count matrix with the “-p -B -t exon -g gene_name” parameters. Raw counts were converted to transcripts per kilobase million and corrected using EdgeR (v.3.36.0) (*70*). Principal component analysis was performed using the prcomp function in R (v.4.1.1) (*71*). GSEA was performed using the fgsea package (v.1.20.0) (*72*) and the hallmark gene set in the Human MSigDB Collections (*49*).

### ChIP-seq

FUCCI-expressing MCF-7 cells were seeded in 100-mm dishes (2 300 000 cells/dish) and cultured in 10% FBS containing DMEM for 48 h. The medium was changed to FBS-free phenol red-free DMEM, the cells were cultured for 17 h, the medium was changed, and the cells were cultured for 3 h. After administration of 10 nM HRG for 12 h, the cells were independently treated with CDK4i and DMSO. Cells were washed twice with PBS at the indicated times. Cells were fixed with 1% formaldehyde (Thermo Fisher Scientific) for 5 min. Fragmented DNA was obtained using a SimpleChIP Plus Enzymatic Chromatin IP Kit (Cell Signaling Technology, 9005) and M220 Focused-ultrasonicator (Covaris, Woburn, MA, USA). Immunoprecipitated DNA was obtained using an Auto iDeal ChIP-seq kit for Histones ×24 (Diagenode, Liege, Belgium) on an SX-8G platform (Diagenode). Next, 2 µg anti-H3K27Ac antibody (Diagenode) and 2 µg anti-H3K4me1 antibody (Diagenode) were mixed with 10 µl DiaMag Protein A-coated magnetic beads, and IP reactions with DNA were performed for 12 h. De-crosslinking was conducted using 4 µl of 5 M NaCl and 1 µl of 20 mg/ml proteinase K (Thermo Fisher Scientific) for 4 h at 60 °C. Library construction and RNA-seq were performed using the Illumina sequencing platform (Rhelixa). A sequencing library was generated using a NEBNext Ultra II DNA Library Prep Kit for Illumina (New England Biolabs, 7645). ChIP-seq was performed on the NovaSeq 6000 (Illumina) with 150-bp paired-end reads. H3K27Ac ChIP-seq and H3K4me1 data were processed using the same protocol as for RNA-seq. FastQC (v.0.11.9) was used for quality control. Trimming was performed using Trimmomatic (v.0.39) (*73*) with the following parameters: ILLUMINACLIP, TruSeq2-SE; fa, 2:30:10; LEADING, 30; TRAILING, 30; SLIDINGWINDOW, 4:15; MINLEN, 30; HEADCROP, 2. Reads were mapped using Bowtie2 (v.2.4.4) (*74*) with GRCh38 as the reference genome, and duplicate reads were excluded using Picard (v.2.26.6) (*75*). Bigwig files were created from the generated BAM files using deepTools (v.3.5.1; bamCoverage function) (*76*). Peak regions were identified using HOMER (v.4.11) (*77*) with the “-style histone -L 0 -fdr 0.00005” parameters. All detected peak regions were annotated to their nearest neighbor genes using HOMER annotatePeaks.pl (v.4.11).

### Promoter and enhancer detection

Promoter regions were defined as those in the H3K27Ac peak that did not overlap with the H3K4me1 peak and were within 250-bp of the transcription start site (TSS, at the center of the peak). Enhancer regions were defined as those in the H3K27Ac peak overlapping with the H3K4me1 peak and were more than 250-bp from the TSS. The TSS was obtained from the Ensembl database using BiomaRt (v.2.50.3) (*78*). The bed files containing the promoter and enhancer regions for all time points were merged, and the time-course of the coverage values for H3K27Ac ChIP-seq was calculated using HOMER annotatePeaks.pl (v.4.11). The estimated coverage values were used to detect promoters and enhancers affected by CDK4i. Promoters that gained or lost signal were defined as those with coverage values of |log2 FC| > 0.3 for inhibitor and control conditions. Enhancers that gained or lost signals were defined as those with coverage values changing by |log2 FC| > 0.3 and target gene expression changing by |log2 FC| > 0.2 in the same direction as the change in enhancer activity.

### Motif analysis of ChIP-seq data

Motif analysis was performed using MEME Suite SEA (v.5.4.1) (*79*). The motif files were obtained from the JASPAR 2022 database (JASPAR2022_CORE_vertebrates_nonredundant_v2) (*80*). Peak regions containing motifs were detected using MEME Suite FIMO (v.5.4.1). The analysis was performed using default parameters.

### Transcription factor enrichment analysis

Transcription factor analysis was conducted using DoRothEA (v.1.6.0) (*42*) with the “run_viper” function. We used regulon data with “confidence” levels of A and B. Target genes were selected based on their changes in expression (FDR < 0.01) between the control and inhibitor treatments at each time point.

### Clinical data analysis

We used microarray gene expression data (GEO GSE93204) for biopsy samples from the NeoPalAna clinical trial (NCT01723774) (*50*) to identify differentially expressed genes based on *P*-values adjusted using the Benjamini–Hochberg method to control the false discovery rate in GEO2R.

### Statistical analysis

Statistical analysis, data visualization, and Kolmogorov–Smirnov normality testing were performed in the R software (v.4.2.1; The R Foundation, Vienna, Austria). For normally distributed data, the Student’s *t*-test and Welch’s two-sample *t*-test were used after confirming equal variances using the F-test for comparisons between two groups. Comparisons between more than two groups were performed using one-way analysis of variance followed by the Tukey’s post-hoc test. For non-normally distributed data, a Kruskal–Wallis test or Mann– Whitney U test was used. *P* < 0.05 was considered significant.

## Acknowledgments

We thank all members of the Laboratory for Cell Systems at the Institute for Protein Research for the discussions regarding our research and especially Mr. Kyoichi Ebata for helping to design the cell cycle analysis. We also thank Mr. Hideya Aragaki from Olympus Corporation for providing the cell-tracking software.

## Funding

JSPS KAKENHI (18H04031, MO; 21K15503, ANI)

JST CREST Program (JPMJCR21N3, MO)

Uehara Memorial Foundation (MO)

Kaneko Narita Research Grant from the Protein Research Foundation (ANI)

New Field Development Support Program from the Institute for Protein Research (ANI)

Joint Research Grant from ExCELLS (MO, KA)

## Contributions

Conceptualization: ANI, MO

Methodology: ANI, KM, KA, MO

Validation: ANI

Formal analysis: ANI, KM

Investigation: ANI, KM Resources: ANI, KA, MO

Visualization: ANI, KM

Writing – Original Draft: ANI, KM, MO

Writing – Review & Editing: ANI, KM, KA, MO

Supervision: ANI, MO

Project administration: ANI, MO

Funding acquisition: ANI, KA, MO

## Data availability

The sequence data for RNA-seq and ChIP-seq reported in this paper have been deposited in the DNA Data Bank of Japan with accession number ESUB001718. The code is available at https://github.com/okadalabipr/HER2_Cellcycle. All other data will be made available upon reasonable request.

