## Supplementary figures for "ErbB2/HER2 governs CDK4 inhibitor sensitivity and timing and irreversibility of G1/S transition by altering c-Myc and cyclin D function"

### Fig. S1

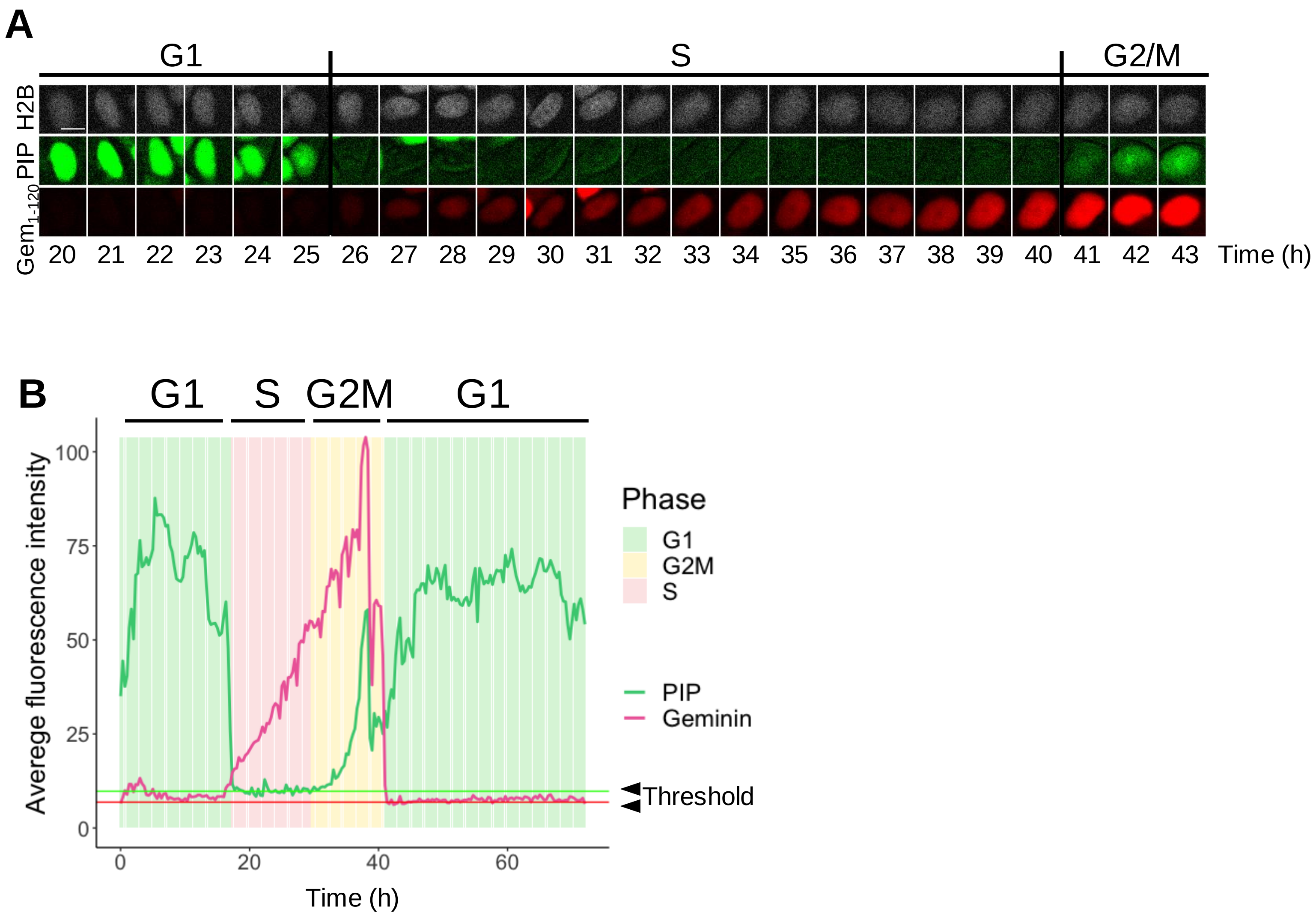


**Fig. S1. Estimation of cell-cycle dynamics through single-cell tracking.** (**A, B**) Data were acquired every 20 min for 72 h after 10 nM HRG stimulation in FUCCI stably expressing MCF-7 cells. (**A**) Representative examples of single-cell tracking with nuclear markers (H2B: gray), G1 and G2M markers (PIP-Cdt1: green), and S and G2M markers (red). (**B**) Cell cycle dynamics estimated below a set threshold from the variation in the average fluorescence intensity of PIP-Cdt1 and Geminin over time. For each cell cycle, green indicates the G1 phase, yellow indicates the G2M phase, and magenta indicates the S phase. The average fluorescence intensity of PIP-Cdt1 and Geminin is indicated by the green and magenta lines, respectively.

### Fig. S2

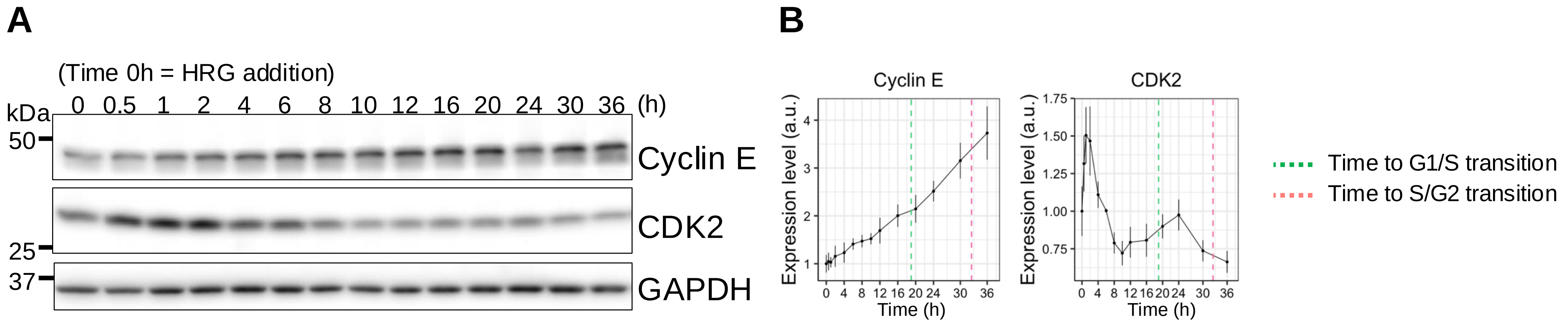


**Fig. S2. Expression dynamics of signaling molecules and cell cycle regulators.** (**A, B**) Expression of proteins regulating the G1/S transition in the ErbB pathway up to 36 h after 10 nM HRG treatment detected using western blotting. (**B**) Expression of each protein normalized relative to that of GAPDH and divided by that of the 0 h sample (0 h = 1). Graphs show means ± SE from three independent replicates.

### Fig. S3

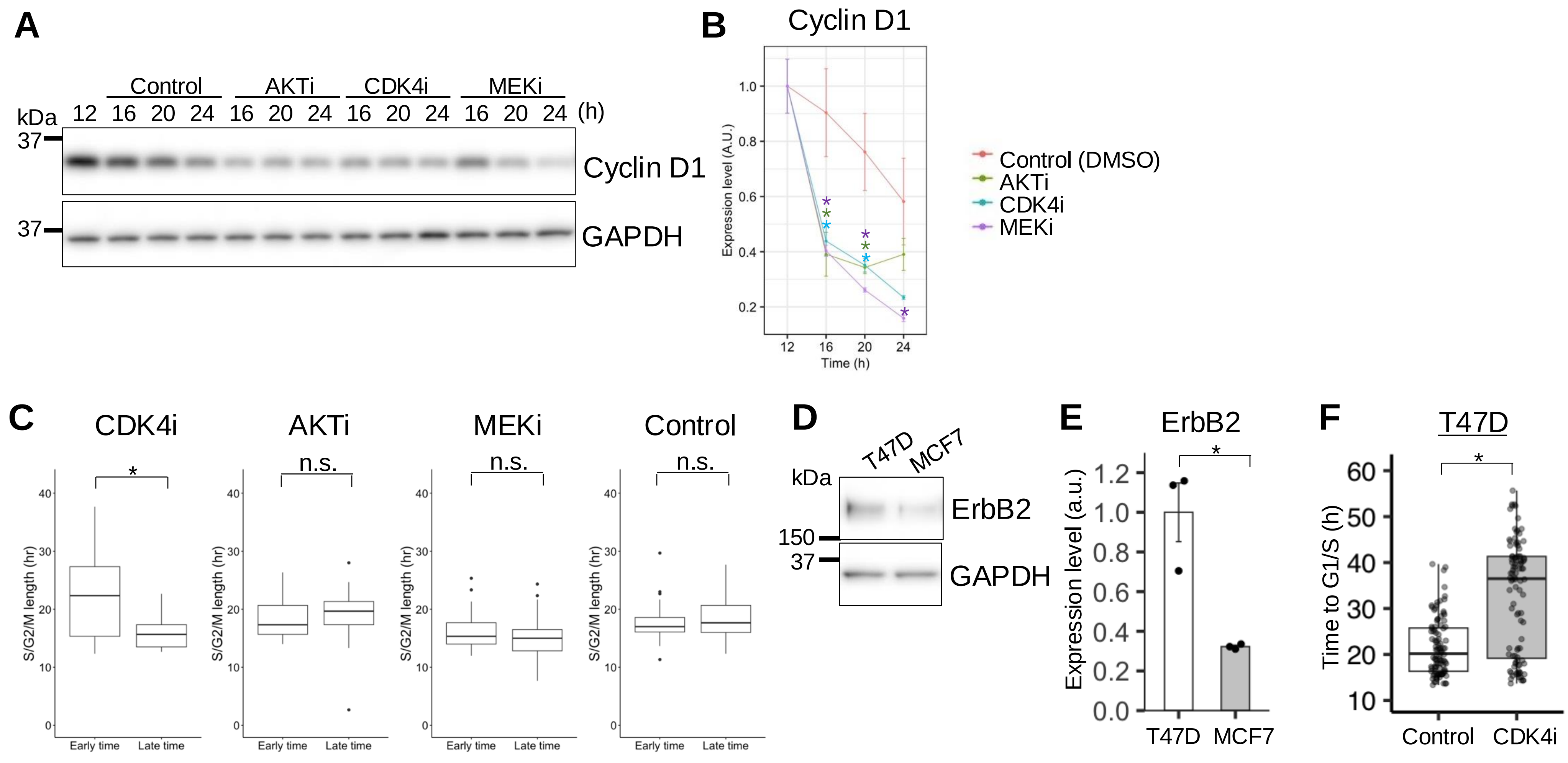


**Fig. S3. Effect of CDK4, AKT, and MEK inhibitors on the expression of cell-cycle regulators and cell-cycle phase distribution.** (**A–C**) FUCCI stably expressing MCF-7 cells were treated with each inhibitor (AKTi: 600 nM; CDK4i: 250 nM; MEKi: 500 nM; DMSO was used as the control) 12 h after 10 nM HRG stimulation. (**A**) Cells were harvested every 4 h until 24 h after 10 nM HRG stimulation and the expression of cyclin D1 and GAPDH was evaluated using western blotting. (**B**) Using the data from (A), the level of each protein was normalized to that of GAPDH and divided by that of the 12 h sample (12 h = 1). Graphs show means ± SE for three independent experiments. (**C**) Images captured every 20 min until 72 h after 10 nM HRG stimulation were used to divide the population into an early and late subpopulation relative to the median time to the G1/S transition and to quantify the length of the S/G2/M phase in each population. CDK4i: $n$ = 27, average time = 15.8 ± 2.8 h (Late time), $n$ = 25, average time = 22.3 ± 7.2 h (Early time); AKTi: $n$ = 50, average time = 18.4 ± 3.9 h (Late time), $n$ = 46, average time = 17.6 ± 2.9 h (Early time); MEKi: $n$ = 38, average time = 15.1 ± 3.6 h (Late time), $n$ = 37, average time = 16.0 ± 3.0 h (Early time); Control: $n$ = 21, average time = 18.9 ± 5.2 h (Late time), $n$ = 21, average time = 18.4 ± 3.6 h (Early time). (**D**) Expression of ErbB2 and GAPDH determined using western blotting. (**E**) ErbB2 level normalized relative to that of GAPDH; $n$ = 3. (**F**) Data from cells that completed the G1/S transition within 72 h. Control and CDK4i; $n$ = 100. In (B), (C), (E), and (F) *$P$ < 0.05, ($t$-test).

Fig. S4

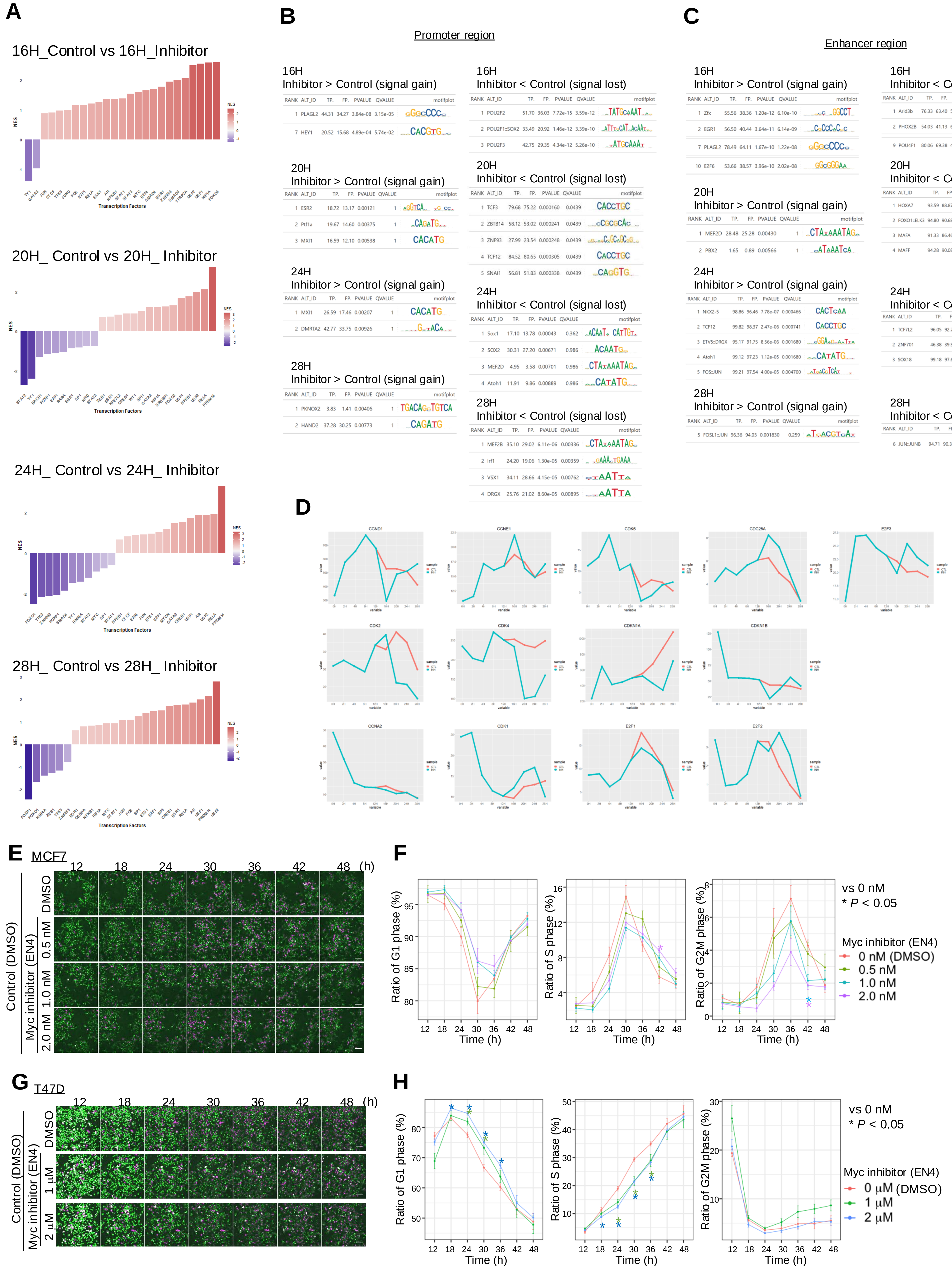
A
16H_Control vs 16H_Inhibitor
20H_ Control vs 20H_ Inhibitor
24H_ Control vs 24H_ Inhibitor
28H_ Control vs 28H_ Inhibitor
Transcription Factors
NES
B
Promoter region
16H
Inhibitor > Control (signal gain)
16H
Inhibitor < Control (signal lost)
20H
Inhibitor > Control (signal gain)
20H
Inhibitor < Control (signal lost)
24H
Inhibitor > Control (signal gain)
24H
Inhibitor < Control (signal lost)
28H
Inhibitor > Control (signal gain)
28H
Inhibitor < Control (signal lost)
C
Enhancer region
16H
Inhibitor > Control (signal gain)
16H
Inhibitor < Control (signal lost)
20H
Inhibitor > Control (signal gain)
20H
Inhibitor < Control (signal lost)
24H
Inhibitor > Control (signal gain)
24H
Inhibitor < Control (signal lost)
28H
Inhibitor > Control (signal gain)
28H
Inhibitor < Control (signal lost)
D
E
MCF7
12
18
24
30
36
42
48
(h)
DMSO
0.5 nM
1.0 nM
2.0 nM
Control (DMSO)
Myc inhibitor (EN4)
F
Ratio of G1 phase (%)
Ratio of S phase (%)
Ratio of G2M phase (%)
Time (h)
vs 0 nM
* P < 0.05
Myc inhibitor (EN4)
0 nM (DMSO)
0.5 nM
1.0 nM
2.0 nM
G
T47D
12
18
24
30
36
42
48
(h)
DMSO
1 μM
2 μM
Control (DMSO)
Myc inhibitor (EN4)
H
Ratio of G1 phase (%)
Ratio of S phase (%)
Ratio of G2M phase (%)
Time (h)
vs 0 nM
* P < 0.05
Myc inhibitor (EN4)
0 μM (DMSO)
1 μM
2 μM

**Fig. S4. Transcriptomic and epigenetic analysis to detect transcriptional regulators associated with CDK4 inhibition.** (**A–C**) FUCCI stably expressing MCF-7 cells were treated with 250 nM CDK4i or DMSO (control) 12 h after 10 nM HRG stimulation. Cells were then harvested every 4 h until 28 h after 10 nM HRG stimulation for RNA-seq and ChIP-seq of H3K27Ac and H3K4me1. (**A**) Transcription factor activity variation inferred using RNA-seq data under control and CDK4i conditions ($\log_2$ FC = CDK4i/control; DoRothEA). Blue and red indicate the decreased and increased activities of transcription factors, respectively. (**B–C**) Promoter (B) and enhancer regions (C) identified based on the H3K27Ac and H3K4me1 ChIP-seq peaks, respectively, and enrichment analysis was performed at each time point for the promoter/enhancer region. The signal under CDK4i conditions (whether gained or lost) was compared to that under control conditions. (**D**) Expression dynamics from 0 to 28 h after 10 nM HRG stimulation for cell cycle-related c-Myc-target genes. (**E–H**) After stimulation with 10 nM HRG for 11.5 h, WT MCF-7 (**E, F**) and WT T47D cells (**G, H**) were treated with 0 (DMSO)/0.5/1.0/2.0 nM EN4 (E, F) or 0 (DMSO)/1/2 mM EN4 (G, H) for 30 min and then with DMSO. Scale bar: 100 µm. The percentage distribution of each cell cycle phase over time was quantified (**F, H**). In (F) and (H), *$P < 0.05$ (with respect to control and each inhibitor condition; *t*-test).

### Fig. S5

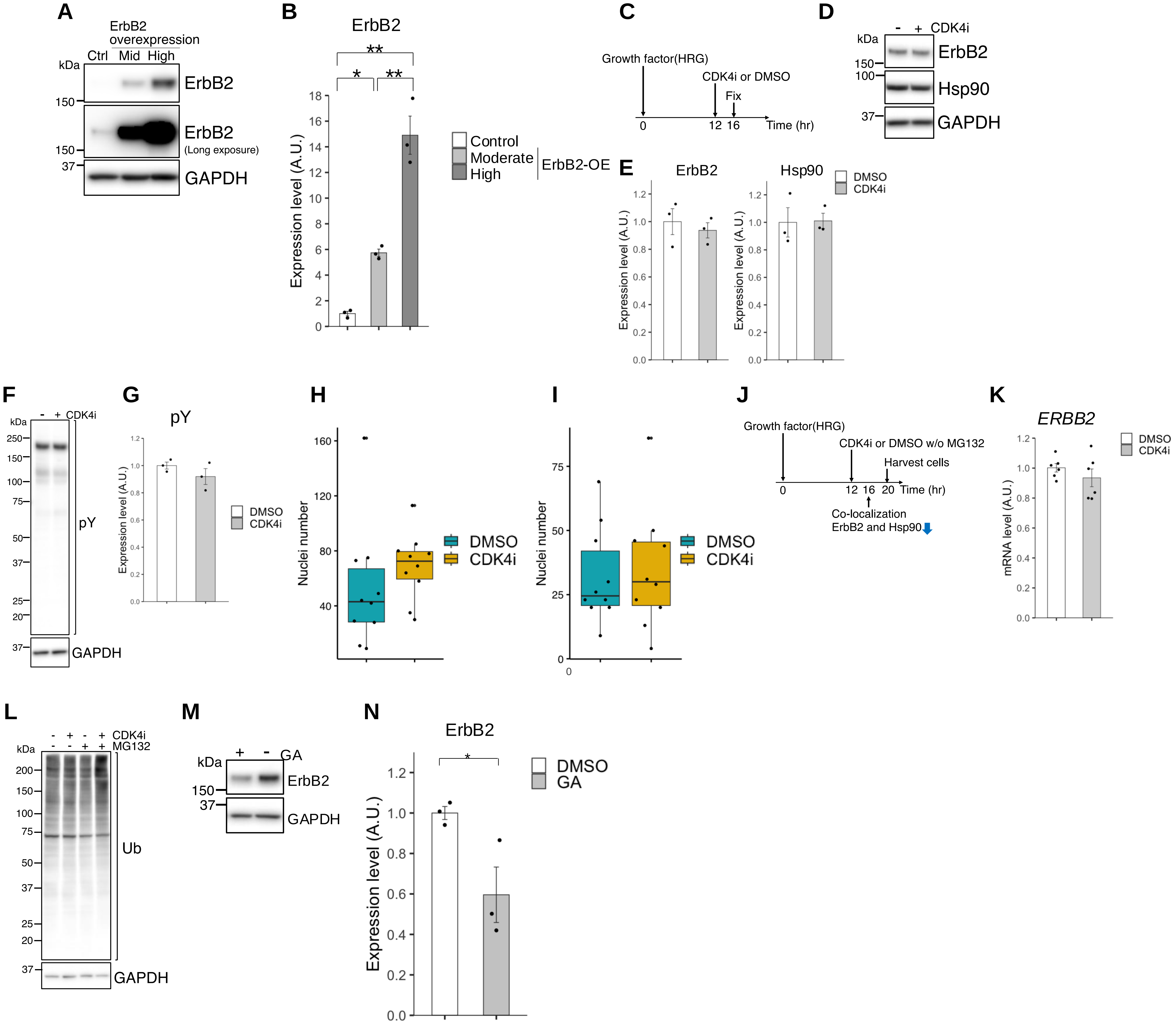


**Fig. S5. G1/S transition requires the stability of ErbB2 protein via Hsp90.** (**A, B**) WT MCF-7 and ErbB2-overexpressing cells (moderate and high levels) were harvested 12 h after 10 nM HRG stimulation. (**A**) Expression of ErbB2 and GAPDH determined using western blotting. (**B**) ErbB2 relative to GAPDH; *n* = 3. (**C–I**) MCF-7 cells overexpressing ErbB2 (moderate levels) were independently treated with CDK4i or DMSO (control) 12 h after stimulation with 10 nM HRG and fixed or collected 4 h later. (**C**) Experimental scheme. (**D**) Expression of ErbB2, Hsp90, and GAPDH determined using western blotting. (**E**) ErbB2 and Hsp90 levels normalized relative to those of GAPDH; *n* = 3. (**F**) Expression of PY100 and GAPDH determined by western blotting. (**G**) PY100 relative to GAPDH, *n* = 3. (**H, I**) Number of nuclei from Fig. 5C (for H) and Fig. 5E (for I). (**J–K**) FUCCI stably expressing MCF-7 cells were treated with CDK4i or DMSO (control) as well as 10 mM MG132 12 h after 10 nM HRG stimulation and collected 8 h later. (**J**) Experimental scheme. (**K**) Comparison of ErbB2 mRNA expression detected using qPCR (*n* = 3 technical replicates with 2 biological replicates). (**L**) Ubiquitin detected using western blotting. (**M, N**) FUCCI stably expressing MCF-7 cells were harvested 8 h after being independently treated with 40 nM geldanamycin (GA) or DMSO (control) 12 h post 10 nM HRG stimulation. (**M**) Expression of ErbB2 and GAPDH determined using western blotting. (**N**) ErbB2 relative to GAPDH; *n* = 3. In (B), **P* < 0.05 (Tukey's multiple comparisons test). In (E), (G), (H), (I), (K), and (N), **P* < 0.05 (*t*-test).

### Fig. S6

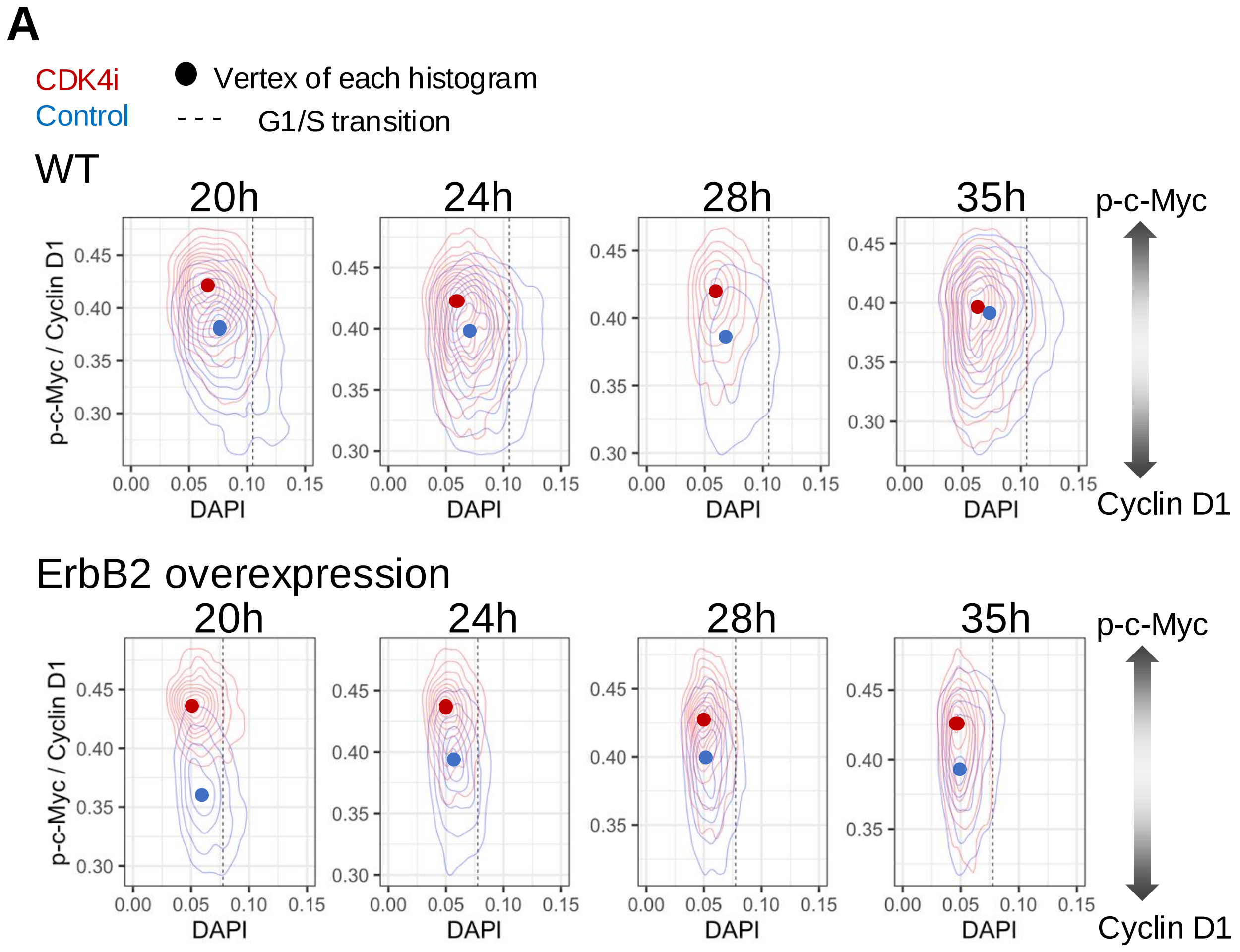


**Fig. S6. p-c-Myc/cyclin D1 ratio is sustained in ErbB2-overexpressing cells compared to control cells.** (**A**) Control and MCF-7 cells overexpressing ErbB2 were individually treated with 250 nM CDK4i or DMSO (control) 12 h after 10 nM HRG stimulation; 20, 24, 28, and 35 h after HRG stimulation, cells were fixed, and p-c-Myc and cyclin D1 were stained for quantification. The vertex of each histogram is indicated by a circle; the G1/S transition estimated from the DAPI staining intensity is indicated by a black dotted line. Blue indicates the control; red indicates the CDK4i condition; $n$ = 3 420 for each condition.

### Table S1. Primers used for real-time PCR

| Gene | Forward primer sequence (5'-3') | Reverse primer sequence (5'-3') |
|---|---|---|
| **CDKN1B** | AGATGTCAAACGTGCGAGTG | TCTCTGCAGTGCTTCTCCAA |
| **ERBB2** | AACTGCACCCACTCCTGTGT | TGATGAGGATCCCAAAGACC |
| **RPL27** | CTGTCGTCAATAAGGATGTCT | CTTGTTCTTGCCTGTCTTGT |
